# Rab7 dependent regulation of goblet cell protein CLCA1 modulates gastrointestinal homeostasis

**DOI:** 10.1101/2023.02.13.528320

**Authors:** Preksha Gaur, Yesheswini Rajendran, Bhagyashree Srivastava, Manasvini Markandey, Vered Fishbain-Yoskovitz, Gayatree Mohapatra, Aamir Suhail, Shikha Chaudhary, Shaifali Tyagi, Subhash C Yadav, Amit K Pandey, Yifat Merbl, Avinash Bajaj, Vineet Ahuja, Chittur V Srikanth

**Affiliations:** Laboratory of gut inflammation and infection biology, Regional Centre for Biotechnology, 3rd milestone Gurgaon Faridabad Expressway, Faridabad 121001, India; Departmnet of Bioscience and Biotechnology, Banasthali Vidyapith, Vanasthali Rd, Aliyabad 304022, India; Department of Gastroenterology, All India Institute of Medical Sciences, Ansari Nagar East, New Delhi, Delhi 110029, India; Department of Immunology, Weizmann Institute of Science, Herzl St 234, Rehovot, Israel; Gene Lay Institute of Immunology and Inflammation, Brigham and Women’s Hospital, Massachusetts General Hospital and Harvard Medical School, Boston, MA, USA; Department of Anatomy, All India Institute of Medical Sciences, Ansari Nagar East, New Delhi, Delhi 110029, India; Vaccine and Infectious Disease Research Center, Translational Health Science and Technology Institute, 3rd milestone Gurgaon Faridabad Expressway, Faridabad 121001, India

## Abstract

Inflammation in ulcerative colitis is typically restricted to the mucosal layer of distal gut. Disrupted mucus barrier coupled with microbial dysbiosis has been reported to occur prior to the onset of inflammation. Here, we show the involvement of vesicular trafficking protein Rab7 in regulating the colonic mucus system. We identified a lowered Rab7 expression in goblet cells of colon during human and murine colitis. *In vivo* Rab7 knocked down mice (Rab7^KD^) displayed a compromised mucus layer, increased microbial permeability and depleted gut microbiota with enhanced susceptibility to dextran sodium-sulfate induced colitis. These abnormalities emerged owing to altered mucus composition, as revealed by mucus proteomics, with increased expression of mucin protease Chloride channel accessory 1 (CLCA1). Mechanistically, Rab7 maintained optimal CLCA1 levels by controlling its lysosomal degradation, a process that was dysregulated during colitis. Overall, our work establishes a role for Rab7 dependent control of CLCA1 secretion required for maintaining mucosal homeostasis.

## Introduction

A close association of gut with diverse microbial antigens demand for well-regulated mechanisms for tolerance and homeostasis. On a cellular level, intestinal epithelium and local immune cells execute diverse regulatory mechanisms to maintain this intestinal homeostasis (Maloy & Powrie, 2011). Goblet cells, a specialized epithelial cell type, participate by secreting mucins that form mucus layer, functioning as the first line of defense against the microbial population (Johansson & Hansson, 2016). Impairment in mucus layer has been observed before intestinal inflammation in ulcerative colitis (UC), one of the two major forms of inflammatory bowel disease (IBD) (Boltin et al., 2013; Johansson et al., 2010). Active UC patients exhibit reduced number of goblet cells and a compromised mucus layer (Singh et al., 2022). This highlights the importance of goblet cell secretory function and mucus layer in UC pathogenesis. Two stratified layers of mucus exist in colon-a loose outer layer permeable to the luminal flora, and a dense adherent sterile inner layer (Atuma et al., 2001). The core structure of mucus is formed of Muc2. Several other known protein constituents of mucus layer are-FCGBP, CLCA1, TFF3 and ZG16 (Hansson & Johansson, 2010). Chloride channel accessory 1 (CLCA1) is a metalloprotease exclusively secreted by goblet cells and forms one of the major non-mucin components of mucus (Nyström et al., 2018). It has Muc2 cleaving properties and thus is involved in intestinal mucus dynamics and homeostasis (Nyström et al., 2019). In view of its ability to protect intestinal epithelium and prevent infections and inflammation, the underlying cause of aspects related to mucus layer alterations during intestinal inflammation needs investigation.

In a recent report focusing on epithelial-immunocyte crosstalk, our group demonstrated a critical role of SENP7, a deSUMOylase in IBD pathophysiology (Suhail et al., 2019). Interestingly, Rab7, among the several other Rab GTPases identified to physically interact with SENP7, was observed in murine model of IBD. Rab7 protein participates in vesicular transport of a cell, influencing protein turnover, secretion, autophagy and intracellular pathogen survival (Deffieu et al., 2021; Gutierrez et al., 2004; Mohapatra et al., 2018; Vanlandingham & Ceresa, 2009; Yap et al., 2018). Each of these functions is arguably essential in intestinal homeostasis, yet these have not been studied in detail in the context of IBD. With all the knowledge provided earlier, it is apparent that Rab proteins may have a role in IBD pathogenesis. But surprisingly, not a lot is known about these proteins and their specific role in IBD. In light of this information, in the present study, we aimed to systematically look at the role of Rab7 in UC pathogenesis. Our data suggests that dysregulated Rab7 during colitis modulates the secretions from goblet cells thus triggering inflammation in the gut.

## Results

### Rab7 expression is reduced during murine and human colitis

To understand the possible role of Rab7 in gut homeostasis and its relevance to colitis we utilized dextran sulphate sodium (DSS) murine model. The animals were fed with 2.5% DSS in their drinking water, their body-weights were regularly monitored. After 7 days of DSS treatment (hereafter DSS mice), they were euthanized and various tissues were harvested for analysis. Compared to control mice which were given normal drinking water, the DSS mice displayed discernible signs of inflammation including reduced body weight, decreased colon length and various histopathological features (as depicted previously by Suhail et. al. 2019). We examined Rab7 expression in inflamed intestine harvested from DSS-colitis mice. A significant reduction of Rab7 protein was seen in tissue lysates of distal colon from DSS mice compared to healthy control (Fig 1A). Immunohistochemistry (IHC) of colon sections revealed ubiquitous presence of Rab7 protein over whole tissue with intensified expression in crypts and other zones of epithelium in healthy mice which got significantly lessened in DSS mice (inset of Fig S1A). To further understand the dynamics of the Rab7 expression change during the course of disease development, DSS treatment was given for different time durations i.e., three days (DSS 3), five days (DSS 5) or seven days (DSS 7) (Fig S1B). Increased duration of DSS treatment resulted in a gradual decrease in mice body weight (Fig S1C) and colon length (Fig S1D) in DSS 7 and DSS 5 mice when compared with DSS 3 mice. DSS 5 and DSS 7 mice also displayed increased splenomegaly (Fig S1E) mirroring severe inflammation seen during human IBD. While whole colonic tissue lysate and crypts (isolated from colons of mice) showed upregulated expression of Rab7 at the early inflammation states (DSS 3-5) (Fig S1F), interestingly, the intestinal epithelial cells (IECs) displayed significant reduction in expression of Rab7 at this stage (day 3) (Fig 1B).

**Fig 1.**
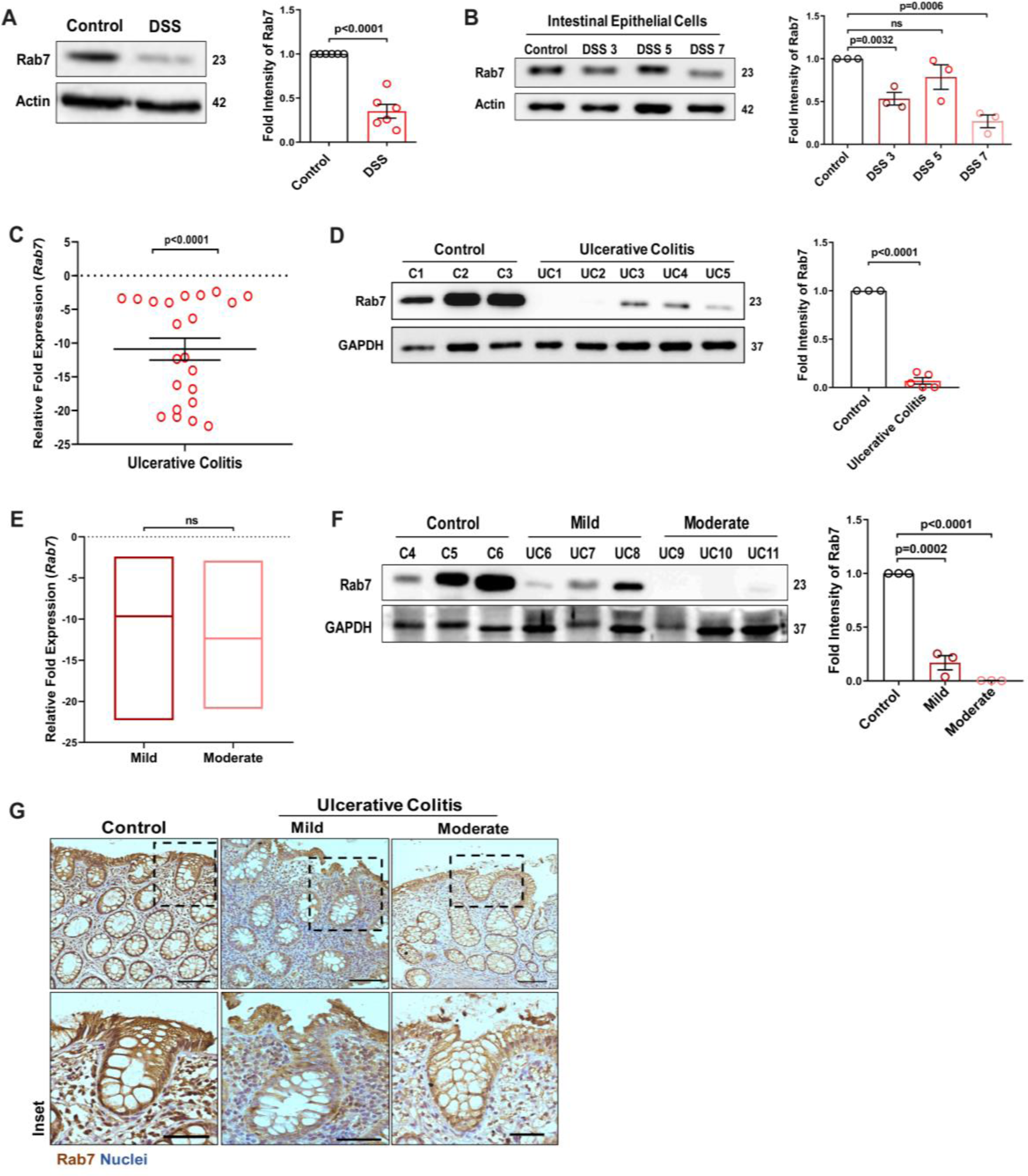
Small GTPase Rab7 shows altered expression during murine and human colitis correlative of disease severity. (A) Rab7 expression analyzed in whole colon tissue of healthy and DSS-treated mice. Graph represents densitometric analysis showing fold intensity of Rab7 expression calculated by normalizing to loading control (β actin). (B) Dynamics of Rab7 expression in intestinal epithelial cells isolated from the intestines of healthy and DSS-treated mice for different time durations. Corresponding graph shows densitometry analysis of Rab7 expression normalized to loading control (β actin). (C) RT-PCR analysis of relative fold expression of *Rab7* gene in human UC patient colonic biopsies (n=22) relative to average control values (n=22). *HPRT* was used for normalization. (D) Immunoblotting of Rab7 protein in human UC (n=5) and control (n=3) biopsy samples. GAPDH was used as loading control. Graph shows densitometry analysis of Rab7 expression relative to control. (E) RT-PCR analysis of *Rab7* gene expression variation with disease severity. (F) Immunoblot showing Rab7 protein expression dynamics of mild (n=3) and moderate (n=3) UC patient samples relative to controls (n=3). Graph represents densitometric analysis showing fold intensity of Rab7 expression calculated by normalizing to loading control (GAPDH). (G) Immunohistochemistry images of colon biopsies from control (n=3), UC mild (n=3) and UC moderate (n=3) patients stained for Rab7 (brown color) (Scale bar=100µm). Inset shows zoomed areas of the image (Scale bar=50µm). Each dot represents (A, B) one mouse or (C, D and F) one human. Error bars represent mean+SEM. Statistical analysis by Student’s t test. ns=non-significant. See also Fig S1 and Table S1.

To investigate if these observations are relevant to Human IBD, we carried out a systematic analysis of Rab7 in human UC endoscopic samples. Details of the patient’s clinical parameters are summarized in Table S1. A total of 65 human samples were utilized for different experiments. The control biopsies in our investigation were acquired from non-IBD patients. qRT-PCR of 22 colonoscopy tissue samples each of UC and controls revealed ∼ -11 fold downregulation of *Rab7* in colitis relative to healthy controls (Fig 1C). The downregulation was also evident at protein levels as revealed by immunoblotting (Fig 1D). In Crohn’s disease (CD), another form of IBD, similar pattern was visible (Fig S1G and Fig S1H).

A recent investigation has indicated elevated Rab7 levels in IBD patients, notably in the colon’s crypt region (Du et al., 2020). This disparity in Rab7 expression could be due to the nature of the samples and the severity of tissue inflammation therein, as also suggested by our findings in DSS-mice dynamics model. To further validate, we evaluated any correlation between Rab7 expression and disease severity. UC patients were grouped into mild (score 2-4), moderate (score 5-6) and remission (score 0-1) based on ulcerative colitis endoscopic index of severity (UCEIS) score. At the transcriptional level, a decrease in Rab7 expression was observed in mild UC patients, with even more reduction in moderate patients compared to controls (Fig 1E). The lowering of expression was correlative to disease severity as noticed by immunoblotting and immunohistochemistry (Fig 1F and Fig 1G). Notably, similar expression alteration of Rab7 was observed in UC remission patients (Fig S1I). Taken together, our data suggests that variations in Rab7 expression levels can be attributed to the extent of gut inflammation.

### Goblet cells of intestine display dominant expression of Rab7

The results of the immunoblotting and qRT-PCR clearly indicate lowering of Rab7 protein levels during inflammation. Although this decrease could be a result of lower cellular expression, a decrease in the number of Rab7-positive cells or an increase in Rab7-negative cells. For a finer understanding, Rab7 was examined for its localization in specific cell types and defined regions of the intestine. Sections of small intestine, proximal and distal colon part of healthy mice were immunostained for Rab7. In line with the earlier data, apart from intestinal crypts, IHC showed Rab7 specific staining in certain portions of the epithelial lining. A closer look revealed that the staining was higher in cells having a vacuolated morphology, a specific characteristic of goblet cells of the epithelium (Fig S2A). Further, colocalizing Rab7 with UEA1, a goblet cell specific marker, confirmed dominant expression in secretory cells of intestine i.e. goblet cells (Fig S2B). Notably, Rab7 was seen in both crypt base and inter crypt goblet cells (inset in Fig S2B). We further examined for additional changes in sub-tissue level Rab7 protein expression pattern during colitis. Interestingly, immunofluorescence-based staining of Rab7 and goblet cells in colonic sections of DSS mice revealed significantly reduced levels of Rab7 distinctively in goblet cells compared to control mice (Fig 2A, Fig 2B and Fig S2C). Similar results were obtained in the case of UC patient colonic biopsy sections with respect to healthy controls (Fig 2C, Fig 2D and Fig. S2D).

**Fig 2.**
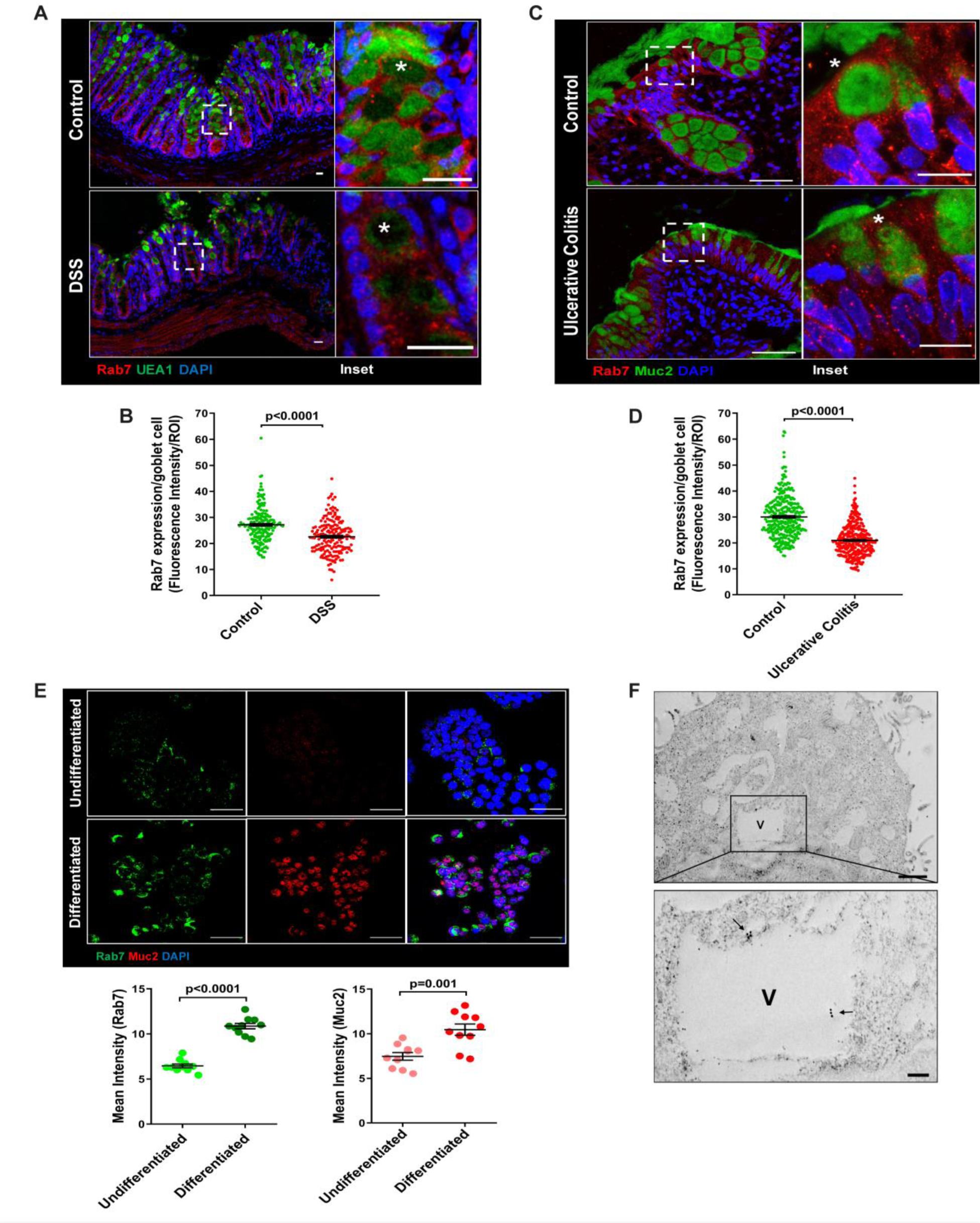
Goblet cells of intestine display dominant expression of Rab7. (A-B) Confocal imaging of healthy and DSS-treated mice (7 days) colon sections showing Rab7 (red) expression within goblet cells (UEA1-FITC) marked with asterisk. Inset shows zoomed areas of the image (scale bar=20µm). Graph shows Rab7 fluorescence intensity within UEA1 positive goblet cells measured within region of interest (ROI). 20 goblet cells were selected randomly from 3 different fields of each mouse (n=3). (C-D) Representative co-immunofluorescence images of human UC and control colon biopsy sections stained for Rab7 (red) and Muc2 (green) for goblet cells (marked with asterisk). (Scale bar=50µm). Inset shows zoomed areas of the image (Scale bar=20µm). Graph shows Rab7 fluorescence intensity within Muc2 positive goblet cells measured within region of interest (ROI). 20 goblet cells were selected randomly from 5 different fields of each sample (n=3). (E) Co-immunofluorescence of Rab7 (green) and Muc2 (red) in undifferentiated and differentiated HT-29 cells (Scale bar=50µm). Graphs show Rab7 and Muc2 fluorescence intensity measured in 10 different fields of three independent experiments each. (F) Rab7 protein visualized in HT29-MTX-E12 cells by immune-EM. Arrows indicate presence of Rab7 on vacuoles of cells (Scale bar=1µm). Inset showed zoomed area of image (Scale bar=200nm). Error bars represent mean+SEM. Statistical analysis by Student’s t test. See also Fig S2.

Goblet cells have a secretory role and as such involve a vigorous and tightly regulated intracellular vesicle trafficking system. Since, Rab7 GTPase is a key regulator of cellular vesicle transport pathway (Guerra & Bucci, 2016), we investigated its possible role in goblet cell function in the context of intestinal inflammation. Pluripotent HT29 cells were cultured in media and condition that is known to induce their differentiation into goblet-like cells (Phillips et al., 1988). Post culturing, the cells showed vacuolated cell morphology along with a loss of expression of Lgr5, stem cell marker, and a concomitant increase in Muc2, a goblet cell specific marker (Fig S2E). Interestingly, compared to undifferentiated HT29 cells, these goblet-like cells showed an increased expression of Rab7 (Fig 2E and Fig S2F). To reconfirm these findings in an alternate cell type, we used HT29-MTX-E12 cells, a differentiated goblet-like cell line. In these cells Rab7 was detectable using gold-labeled antibodies in immune-electron microscopy. Moreover, ultrastructural details (Fig S2G) showed Rab7 protein to be present in the vicinity of secretory vesicles (Fig 2F). Taken together, these data suggest that Rab7 may be involved in regulating the goblet cell function in the intestine.

### Knock-down of Rab7 *in vivo* aggravates DSS-induced colitis in mice

A reduction in Rab7 protein level during colitis, specifically in goblet cells prompted us to further investigate its role in intestinal inflammation. Rab7 knockout mice are embryonic lethal, therefore we developed a transient Rab7 knock-down mice model using an in-house nanogel based oral nucleic acid delivery system (Kawamura et al., 2020; Yavvari et al., 2019). Briefly, mice were fed with either scrambled (C^Scr^) or Rab7 specific siRNA (Rab7^KD^) engineered with nanogel. A sub-group of each category of mice was then fed with DSS (DSS+C^Scr^, DSS+Rab7^KD^) or left untreated (C). Schematic representation that summarizes the different treatments to mice is presented (Fig 3A). Interestingly, only 4 days of DSS administration was sufficient to generate severe inflammation in Rab7^KD^ mice which was atypical for DSS-colitis model. This was evident by reduced body weight (Fig 3B), diarrhea and rectal bleeding (Fig 3C). These mice were dissected and relevant organs were harvested for further analysis. Intestinal epithelial cells were isolated from colon to check for the expression of Rab7 after siRNA treatment through western blot. Almost 50% of protein expression was reduced in Rab7^KD^ mice groups, thus confirming successful knock-down (Fig 3F). No significant alterations in Rab7 expression were visible in other organs such as spleen, MLN and liver proving no off-targeting of nucleic acid by nanogel (Fig S3A). DSS and DSS+C^Scr^ mice showed a significant reduction in colon length with inflamed appearance in comparison to C, C^Scr^ and Rab7^KD^ groups (Fig 3D). Notably, the features were more severe in the case of DSS+Rab7^KD^ group in comparison to DSS+C^Scr^ mice. In line with this we observed splenomegaly in DSS+Rab7^KD^ group depicting higher inflammation (Fig 3E). H&E staining of distal colon of DSS+ Rab7^KD^ mice showed exacerbated signs of inflammation such as immune cell infiltration, colonic wall thickening, goblet cell and crypt loss and epithelial erosion (Fig 3G). While many of these signs were evident in DSS+C^Scr^ mice, in DSS+Rab7^KD^ group the intensity was higher as can be seen in the histopathology score plot (Fig 3H). Mucus scrapings from the colons of different groups of mice were examined for the presence of TNFα, the key proinflammatory cytokine. ELISA revealed a significant upregulation of TNFα in DSS+C^Scr^ mice, which was further higher in DSS+Rab7^KD^ mice (Fig 3I). These data led us to conclude that downregulation of Rab7 heightens DSS induced colitis in mice, thus hinting towards its functional importance during colitis.

**Fig 3.**
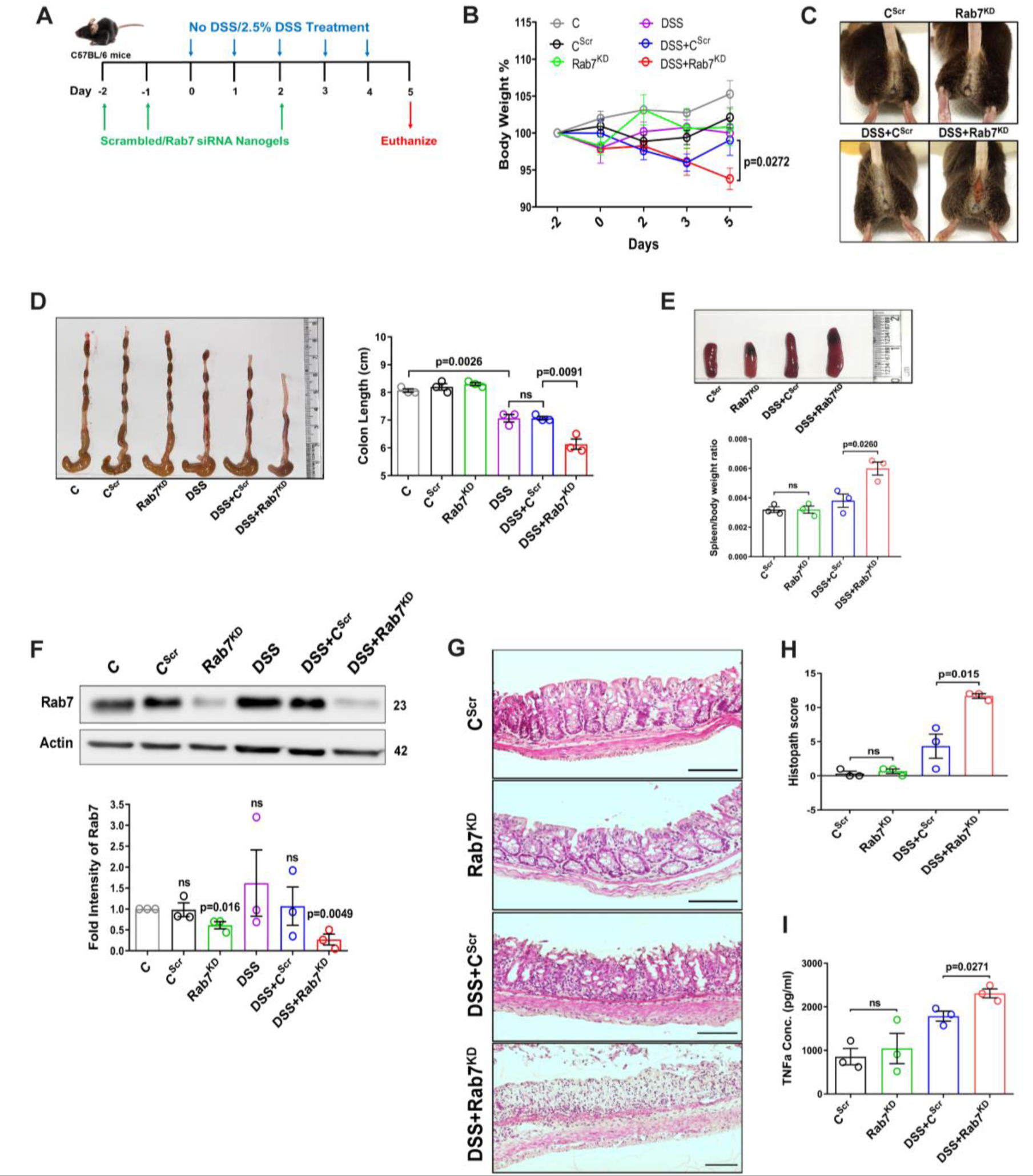
Downregulation of Rab7 aggravates inflammation upon external trigger. (A) Schematic representation of the experimental plan for knock-down of Rab7 in C57Bl/6 mice showing different treatments (n=3 mice per group). (B) Graph showing body weight percent. (C) Representative photographs demonstrating rectal bleeding in mice. (D) Gross morphology of colon and caeca. Graph shows colon length quantification. (E) Representative spleens of different treatment mice groups. Graph showing spleen to body weight ratio. (F) Rab7 protein expression in isolated intestinal epithelial cells from mice colon. Graph represents densitometric analysis showing fold intensity of Rab7 expression calculated by normalizing to loading control (β actin). Significance value of each group is relative to untreated (C) group. (G-H) Hematoxylin and eosin staining of distal colon sections with histopathology scores showing increased characteristics of inflammation. (I) ELISA of TNFα from mucosal extracts of mice colon. Each dot represents one mouse. Error bars represent mean+SEM. Statistical analysis by (B) Two-way Anova or Student’s t test. ns=non-significant. See also Fig S3.

### Depletion of Rab7 in intestinal epithelium modulates mucus layer permeability and gut microbiota

Goblet cells primarily function to secrete mucins which release to form a mucus layer protective for the mucosal surface. Dysfunction of goblet cells leading to reduced mucus layer secretion and decreased barrier function is a frequent abnormality reported in UC (Gersemann et al., 2009). We observed a conspicuously higher expression of Rab7 in goblet cells in steady state, while reduction during colitis as seen in Fig 2B and Fig 2D. Based on these data, we hypothesized that Rab7 perturbation would impact goblet cell function and mucus secretion. To test this, histological examination of the colon from various groups described above was carried out through Alcian-blue staining for mucins. DSS-treated mice groups showed a significantly decreased staining indicating reduced goblet cell number. Notably, this phenotype was observed in Rab7^KD^ mice colon also, although the data was not significant (Fig S3B and Fig S3C). For the analysis of the mucus, thickness of the inner mucus layer (IML) was measured in Alcian blue-stained colon specimens. As expected, the decrease in goblet cell number was accompanied by thinning of the mucus layer in both the DSS-treated mice groups (Fig 4A). Surprisingly, in Rab7^KD^ mice IML was significantly much wider and seemed to be less dense showing a lighter stain of alcian blue (Fig 4B). This led us to expect alterations in the permeability of the mucus layer. We speculated that depletion of or lack of Rab7 may be resulting in compromised mucus properties. To test this, initially a Rab7 knockout HEK cell line (hereafter HEK Rab7^-/-^) was generated. Knockout in intestinal epithelial cells was attempted but we were unable to obtain a line without Rab7 expression. HEK Rab7^-/-^or HEK WT cells were transfected with pSNMUC2-MG plasmid (a kind gift from Professor Gunnar C Hansson, University of Gothenburg). This plasmid expresses human Muc2 N-terminal domains fused with GFP (Godl et al., 2002). Surprisingly, the Muc2 protein was seen to be not forming a theca and was scattered over cell cytoplasm as compared to wildtype cells corroborating the above *in vivo* data (Fig S3D).

**Fig 4.**
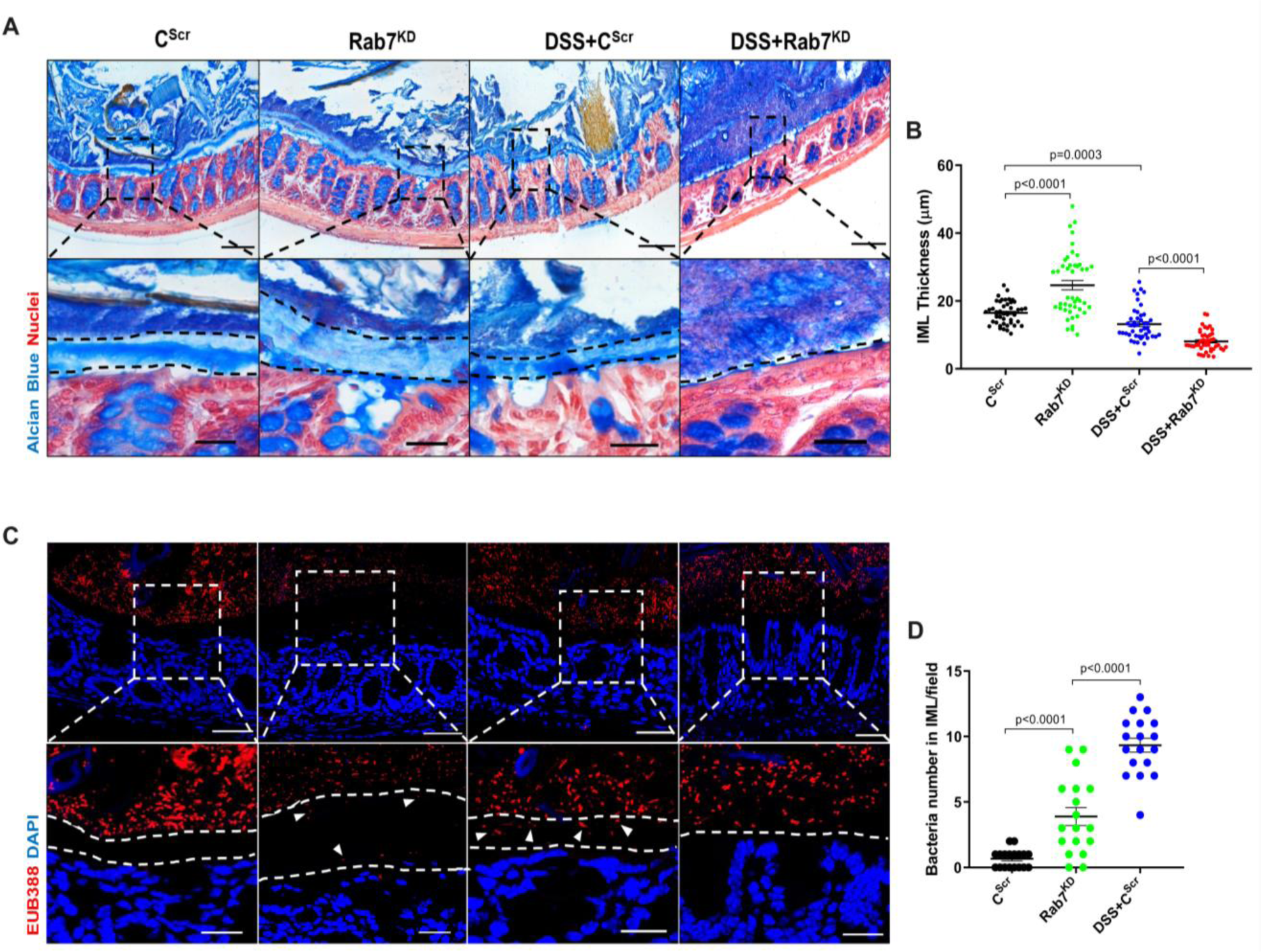
Rab7 downregulation in intestinal epithelium modulates mucus layer thickness and permeability. (A-B) Alcian blue staining in mice distal colon sections displaying IML (blue) marked with dotted lines and nuclei (red). Thickness of IML was measured using Image J software (15 measurements per section of each mice). Scale bar=100µm and 20µm (inset). (C-D) Representative images of FISH staining for bacteria detection in mucus layer using general bacterial probe EUB338-Alexa Fluor 647 (red). Arrow heads demarcate presence of bacteria in IML (dotted lines). Graph shows bacteria count detected in IML (6 regions per section of each mouse). Scale bar=100µm and 20µm (inset). Error bars represent mean+SEM. Statistical analysis by Student’s t test.

To check for any alterations in the permeability of the IML of Rab7^KD^ mice, we analyzed the colon sections, using published protocol, for bacterial presence in the mucus layer by fluorescence-in situ hybridization (FISH) involving a universal bacterial probe EUB388. In control mice as anticipated, bacteria were detected in the outer mucus layer but not in inner layer (Fig 4C). On the contrary, in Rab7^KD^ mice, in spite mucus layer being wider several bacteria were detected in the inner mucus layer verifying increased permeability (Fig 4D). DSS+C^Scr^ mice showed a dramatic increase in number of bacteria penetrating the inner layer. As, there was complete invasion of bacteria to epithelium surface in case of DSS+Rab7^KD^ treated mice, quantification of the number of penetrating bacteria was unworkable. These data clearly indicated towards the involvement of Rab7 in modulating the mucus secreting function of goblet cells which further alters the mucus layer characteristics leading to bacterial breach and generation of inflammation.

Besides mucus barrier integrity, intestinal microbiota plays an important role in gut inflammation (Al Bander et al., 2020). We analyzed the microbial community in stool samples of Rab7^KD^ and DSS+C^Scr^ compared to C^Scr^ mice. Our main motive here was to specifically examine alterations in microbial diversity/abundance in Rab7^KD^ mice relative to DSS+C^Scr^ and C^Scr^ mice. Fecal samples were collected at day 5 of DSS treatment in Rab7^KD^ mice model, particularly since these mice display significant inflammation by day 5 due to Rab7-deficiency. As anticipated, Shannon index plot showed decreased microbial diversity richness and evenness in DSS+C^Scr^ mice groups (µ_median_=3.67). This phenotype was more severe in Rab7^KD^ mice (µ_median_=3.46) (Fig 5A). Further, principal-coordinates analysis (PCoA) indicated that each of the three groups assumed a discrete cluster (Fig 5B). Similar divergence among the DSS+C^Scr^ and C^Scr^ were seen in non-metric multidimensional scaling (NMDS) plots (Fig S4A). While, each group displayed presence of some unique Operational Taxonomic Units (OTUs), some shared OTUs were also notable (Fig 5C). Beta diversity index heatmap revealed differences in the microbial diversity among different experimental mice groups (Fig S4B). The map clearly indicated a shift in microbial diversity in Rab7^KD^ mice such that they appeared to occupy an intermediate position between C^Scr^ and DSS+C^Scr^ mice. Substantial differences in the relative abundance of many microbes were found (Fig 5D). At the phylum level, bacteroidetes, showed an increase in abundance, whereas firmicutes were significantly less represented in Rab7^KD^ and DSS+C^Scr^ mice groups (Fig 5E). The other analyzed phyla did not show any significant changes. Firmicutes are known to get less abundant in IBD (Frank et al., 2007), (Manichanh, 2006). Class level analysis revealed that the Firmicutes reduced were clostridia, while the abundance of Bacteroidia was more among Bacteroidetes (Fig 5F). Abundance of several other microbes getting altered could be appreciated at order, family, genus and species level (Fig S4C). Genus such as *Lactobacillus* and *Parabacteroidetes* were found to be significantly altered in Rab7^KD^ mice in comparison to C^Scr^ (Fig 5G). Intriguingly, compared to C^Scr^ mice, Rab7^KD^ and DSS+C^Scr^ mice showed a decrease in the abundance of a particular category of bacterial species (Fig S4D, red box). The majority of these contained probiotic microorganisms. These include *Bifidobacterium longum* and *Lactobacillus helveticus*, both of which have been thoroughly researched in relation to colitis (Ho et al., 2022; Tamaki et al., 2016; Zhang et al., 2021). All these data demonstrated aberrant composition of gut microbiota of Rab7^KD^ mice with more similarity with DSS mice inflamed gut microbiota.

**Fig 5.**
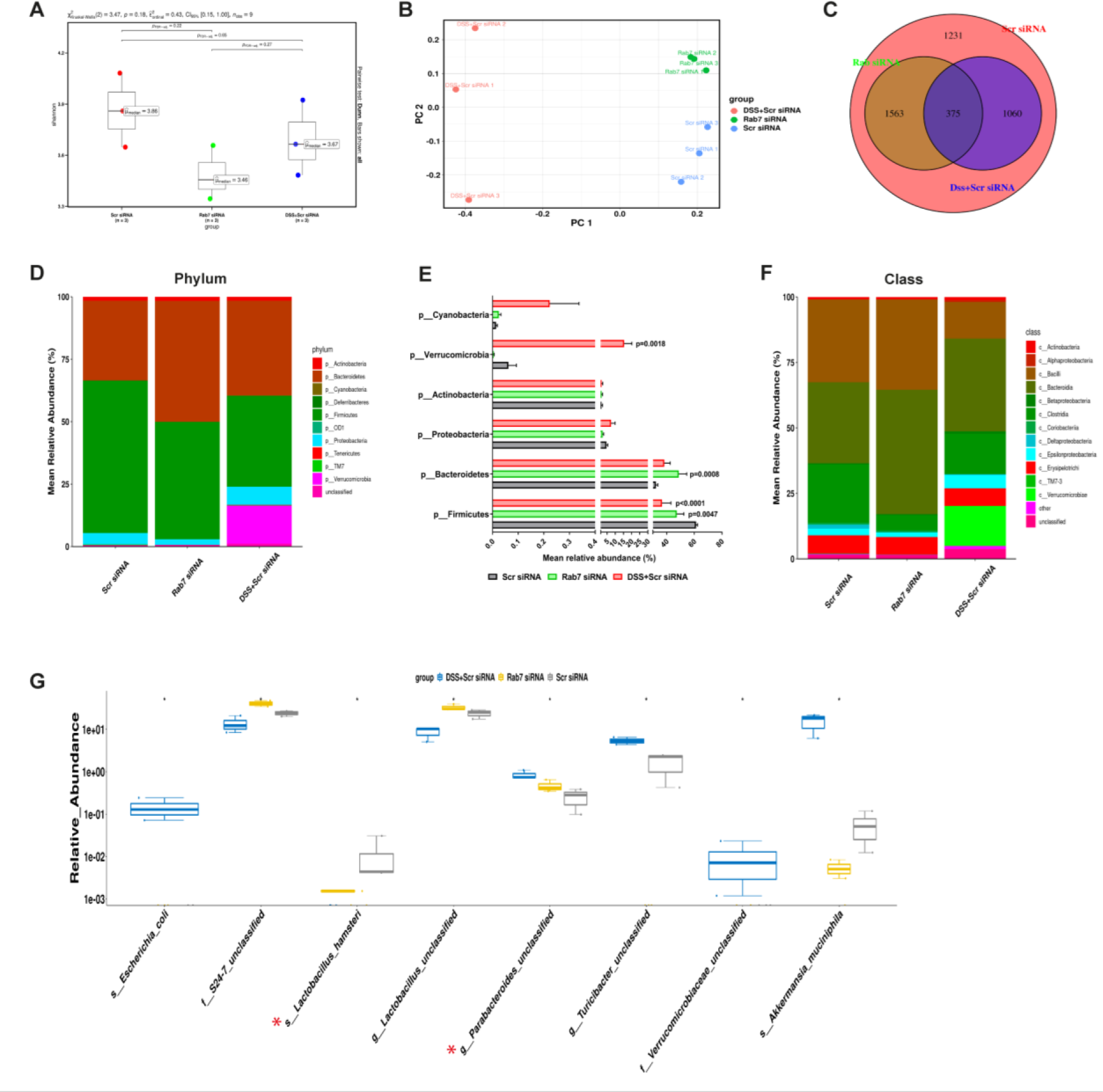
Gut microbiota in Rab7 knockdown mice is altered alike DSS-colitis mice. Microbial composition of C^Scr^ (Scr siRNA), Rab7^KD^ (Rab7 siRNA) and DSS+C^Scr^ (DSS+Scr siRNA) was analyzed by 16S metagenomic profiling. (A) Alpha diversity quantified as Shannon index. Significance was calculated using the Kruskal-Wallis test followed by the improved Benjamini-Hochberg procedure from false discovery rate (FDR) correction. (B) Principle Coordinates (PCoA) plot calculated from distance matrices obtained from Bray-Curtis. (C) Venn diagram representation of shared and unique OTUs between groups. (D-F) Mean relative abundance of each taxon. Relative abundance of top 10 Phylum (D) and Class (F) of each group is depicted in the figure. Each bar represents the mean of the merged OTUs from three mice. Taxonomic lineages not included in top 10 were collapsed as “Others”, while the ones which have not been classified has been placed under the category of “unclassified”. Relative abundance (in %) of some important taxa showing significant differences between experimental groups (E). (G) Plot showing relative abundance of 8 different taxa, identified with Kruskal-wallis test. Each dot represents one mouse. See also Fig S4.

### Rab7 perturbation impacts mucus composition in colon

We next sought to investigate in details the alterations in the colonic mucus composition in Rab7 perturbed mice which might influence penetrability and gut microbiota. The secreted mucus is composed of a mixture of proteins, including those contributing to mucus gel architecture, antimicrobial peptides and regulatory processes. We went on to investigate any modifications in the total amount or composition of the secreted mucus of Rab7^KD^ mice. Mice colon was harvested and secreted mucus was isolated by gentle scrapping using a rubber policeman (Fig 6A) and run on a mucin gel stained with alcian blue dye. No significant change was observed in the total mucin content between any of the groups (Fig 6B). We speculated that there may be changes in specific components of mucus. For identification of proteins in the secreted mucus, isolated mucus samples were subjected to reducing buffer and separated on SDS-PAGE followed by high resolution tandem mass spectrometry (mucus proteomics) (Fig 6C). Raw data was analyzed in MaxQuant software (v1.6.0.16). Masses were searched against the mouse UniProt proteome database and additional databases such as miceMucinDB and VerSeDa. 522 proteins were detected among different mice groups (Fig 6D) of which 288 were common in all along with some unique proteins in each group as is evident through the venn diagram (Fig 6E). Principal component analysis (PCA) showed proper grouping of the biological replicates from different experimental groups (Fig S5A). Label free quantification identified approximately 500 differentially expressed proteins among the samples (Fig 6F). Significantly altered proteins were analyzed with FDR (Fig 6G). Proteins observed to be differentially expressed in Rab7^KD^ mice in comparison to C^Scr^ mice or DSS+Rab7^KD^ mice and DSS+C^Scr^ mice are represented in respective volcano plots (Fig 6H and Fig 6I). These proteins were part of certain pathways over represented in Gene ontology (GO) analysis like extracellular exosome biogenesis, vesicle localization, calcium-dependent phospholipase A2 activity based on biological processes (Fig S5B) and molecular function (Fig S5C). All the proteins which displayed significant differential expression were localized in different subcellular compartments (Fig S5D and Fig S5E). Some cytoplasmic and nuclear proteins were also represented in this which may be debris or exudates from lysed cells. Certain secretory proteins such as Chloride channel accessory 1 (CLCA1), Apolipoprotein AI (APOA1) and Transferrin (Tf) caught our interest as they were seen to be upregulated in Rab7^KD^ mice group when compared with the C^Scr^ (highlighted in red in Fig S5D and Fig S5E). As, transferrin or Tf is an iron binding glycoprotein secreted from the liver, their increased amount in mucus indirectly hinted towards the decrease in transferrin receptors (TFRC) in the intestinal cells which uptake them. We therefore examined the expression level of *CLCA1*, *APOA1* and *TFRC* in colon tissue through qRT-PCR analysis. While *TFRC* showed down expression in DSS+Rab7^KD^ mice, no significant change was observed for *CLCA1* and *APOA1* (Fig S5F, Fig S5G and Fig S5H). In case of *CLCA1*, several folds downregulation at RNA level in Rab7^KD^ and DSS+Rab7^KD^ mice was seen, although the changes were non-significant. CLCA1 is a protease exclusively secreted by goblet cells and forms one of the major mucus components. It has recently been demonstrated that CLCA1 helps in maintaining the dynamics of mucus layer in colon by processing Muc2. The CAT/Cys and VWA domain containing part of CLCA1 protein (53 kDa) has maximum Muc2 cleaving property. We went on to check for the expression of CLCA1 at protein levels in the mucus of Rab7^KD^ mice. Remarkably, CLCA1 (53 kDa) was found to be highly upregulated in mucus samples of Rab7^KD^ followed by DSS-treated mice compared to controls (Fig 7A). CLCA1 could thus be an interesting and probable candidate to justify modifications in mucus layer with its increased secretion in mucus upon Rab7 knock-down in mice.

**Fig 6.**
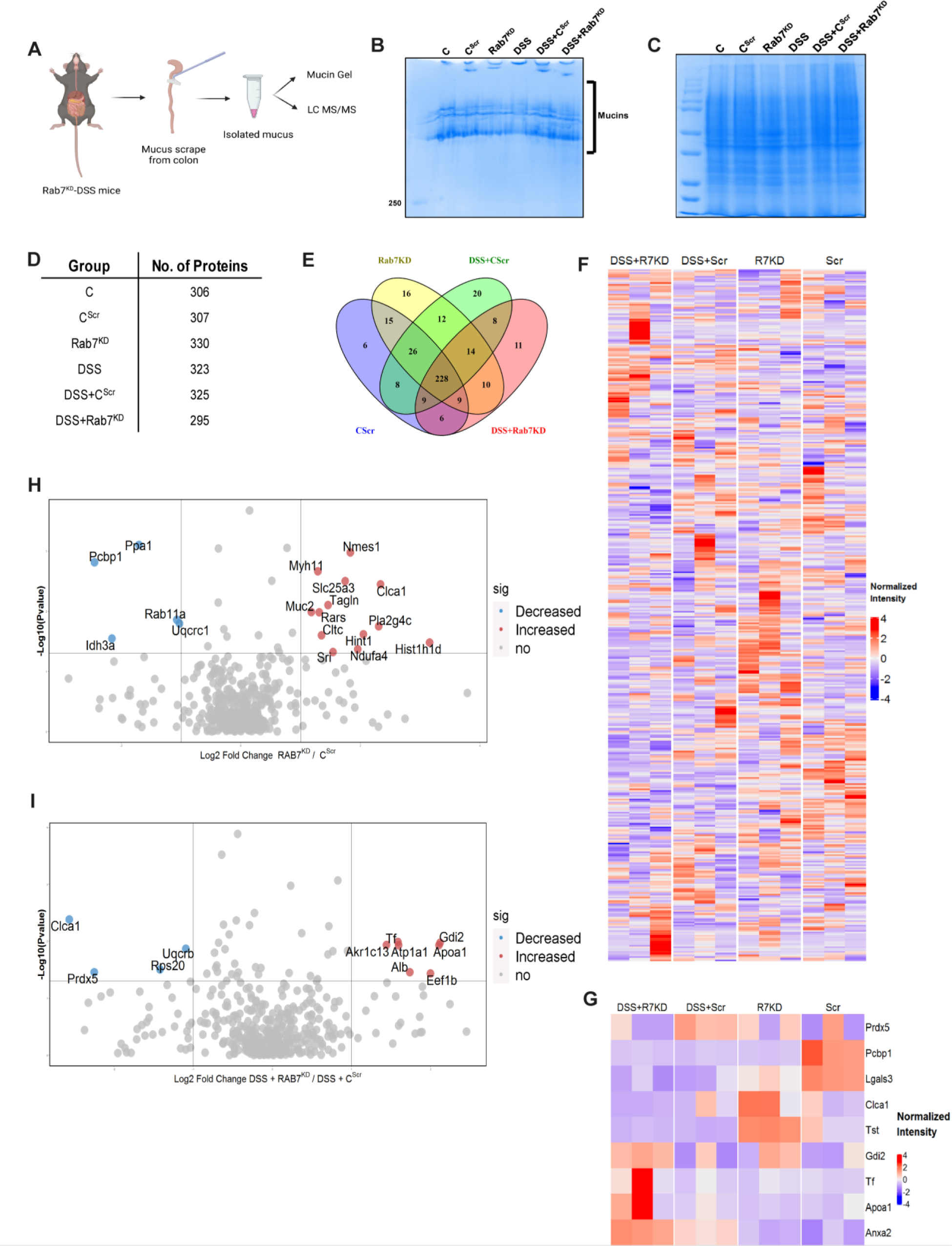
Rab7 perturbation alters mucus composition of colon. (A) Schematic representing steps for mucus isolation from mice colon followed by sample preparation for total mucin measurement through mucin gels and mucus layer composition analysis using mass spectrometry (Created with BioRender.com). (B) Mucin gel showing the amount of total mucins (blue) stained with alcian blue dye.(C-E) Coomassie staining of mucus samples isolated from different experimental groups. Table listing the total number of proteins identified in the mucus samples through mass spectrometry and venn diagram displaying the number of individual proteins common and unique in different samples. (F-G) Heat maps of the proteins differentially expressed without FDR (F) and with FDR (G). (H-I) Volcano plots of proteome in mucus samples of different mice groups showing differential expression of proteins in Rab7^KD^ mice verses C^Scr^ (H) and DSS+ Rab7^KD^ mice verses DSS control group (I). See also Fig S5.

**Fig 7.**
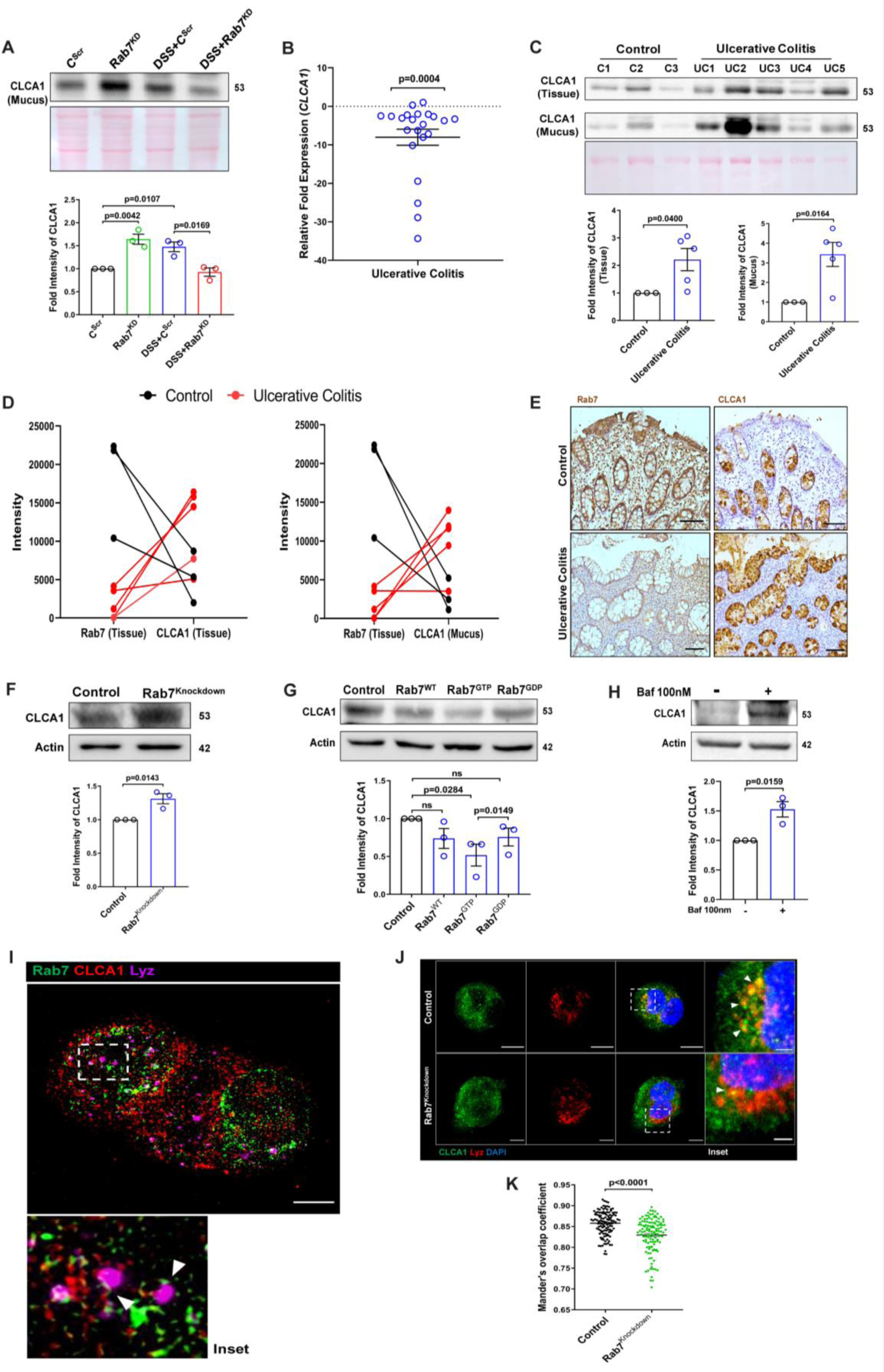
Rab7 mediates CLCA1 degradation via lysosomal pathway in goblet cells. (A) CLCA1 expression in mucus samples of Rab7^KD^-DSS mice. Graph represents densitometric analysis showing fold intensity of CLCA1 expression. (B) RT-PCR analysis of relative fold expression of *CLCA1* gene in human UC (n=22) patient colonic biopsies relative to average control (n=22) values. *HPRT* was used for normalization. (C) Immunoblotting of CLCA1 protein in human UC (n=5) and control (n=3) mucus and biopsy samples. Graphs represent densitometric analysis showing fold intensity of CLCA1 expression in mucus and tissue samples calculated by normalizing to loading control (GAPDH shown in Fig1F). (D) Correlation graph showing expression of Rab7 and CLCA1 in mucus and tissue samples of human UC patients relative of controls plotted using fold change from immunoblots. (E) Representative immunohistochemistry images of Rab7 and CLCA1 staining (brown color) in human UC patient (n=3) and control biopsy (n=3) sections and cell nuclei (blue color) (Scale bar=100µm). (F) CLCA1 protein expression in HT29-MTX-E12 cells transfected with either scrambled siRNA (control) or Rab7 siRNA (Rab7^Knockdown^). Graph represents densitometric analysis showing fold intensity of CLCA1 expression calculated by normalizing to loading control (β actin). (G) Immunoblot showing CLCA1 protein expression change in HT29-MTX-E12 cells overexpressed with EGFP empty vector (control), Rab7-GFP (Rab7^WT^), Rab7-GFP GTP locked form (Rab7^GTP^) and Rab7-GFP GDP locked form (Rab7^GDP^). Graph represents densitometric analysis showing fold intensity of CLCA1 expression calculated by normalizing to loading control (β actin). (H) Immunoblot showing CLCA1 protein expression after treatment of bafilomycin in HT29-MTX-E12 cells. Graph represents densitometric analysis showing fold intensity of CLCA1 expression calculated by normalizing to loading control (β actin). (I) Representative image of structured illumination microscopy showing images of HT29-MTX-E12 cells transfected with Rab7^GTP^ (green) and stained with CLCA1 (Red) using anti-CLCA1 antibody and lysosomes with LysoTracker Red DND-99 (Magenta) (Scale bar=5µm). Inset shows zoomed areas of colocalization marked with arrows. (J-K) Representative confocal images of HT29-MTX-E12 cells transfected with either scrambled siRNA (control) or Rab7 siRNA (Rab7^Knockdown^). Cells are stained with CLCA1 (green) using anti-CLCA1 antibody and lysosomes with LysoTracker Red DND-99 (red). Graph shows quantitation of colocalization between CLCA1 and lysosomes from images (n=120) using Mander’s overlap coefficient. (Scale bar=100µm). Inset shows zoomed areas of the image with colocalization puncta (yellow) marked with arrows (Scale bar=50µm). Each dot represents (A) one mouse or (B, C and E) one human or (F, G and H) one independent experiment. Error bars represent mean+SEM. Statistical analysis by (C) Welch’s t test or (A, B, F, G, H and K) Student’s t test. ns=non-significant. See also Fig S6.

### Altered Rab7 influences CLCA1 expression in goblet cells

We investigated the expression alteration of CLCA1 in human UC. Earlier, van der Post et al. 2019 reported CLCA1 to be getting downregulated in active UC patients. In line with this, qRT-PCR analysis of *CLCA1* in 22 UC and control biopsy samples each revealed significant downregulation in UC relative to healthy controls (Fig 7B). Next, expression of CLCA1 protein was investigated in tissue and secreted mucus from UC patients (same as used for analyzing Rab7 expression in Fig 1D). To achieve that, human colon biopsies were treated with N-Acetyl cysteine, a mucolytic agent, to isolate secreted mucus (Fig S6A). The mucus samples obtained this way and the remaining tissues were processed separately for immunoblotting. Interestingly, similar to the Rab7 knock-down mice, a higher expression of CLCA1 was seen in these human UC samples compared to healthy controls both in tissue and secreted mucus (Fig 7C). Moreover, there was an inverse correlation between expressions of Rab7 and CLCA1 both in tissue and secreted mucus of human colon biopsies (Fig 7D). Also, the immunostaining of Rab7 and CLCA1 in human UC and control colon sections revealed higher expression of CLCA1 in crypts and goblet cells of UC compared to healthy controls, opposite to the pattern seen for Rab7 (Fig 7E). These data depicted that CLCA1 protein expression is increased in tissue resulting in higher secretion into mucus during colitis.

To examine the mechanism of possible Rab7 mediated control of CLCA1 secretion, we used HT29-MTX-E12, cultured goblet cells. siRNA mediated knock-down of Rab7 was carried out in these cells followed by analysis of CLCA1 protein expression. We observed an increase in CLCA1 expression in the Rab7 knocked down cells (Fig 7F). Further, overexpression of Rab7-GTP locked form (active Rab7) lowered CLCA1 in cells (Fig 7G). All this data led us to anticipate the involvement of Rab7 in regulating the CLCA1 expression in the goblet cells of intestine.

Since currently there are no reports addressing CLCA1 protein stability in goblet cells, we carried out experiments to delineate this further. HT29-MTX-E12 cells were treated with MG132 (proteasome inhibitor) and bafilomycin (lysosome inhibitor) and CLCA1 expression was checked. CLCA1 showed rescue upon bafilomycin treatment confirming that it gets degraded via lysosomal pathway (Fig 7H). No significant change was seen in CLCA1 expression upon proteasomal pathway inhibition (Fig S6C). We speculated a role for Rab7 in regulation of CLCA1 protein turnover via lysosomal degradation pathway, since Rab7 GTPase is known to facilitate fusion of late endosomes to lysosomes for degradation. Structured illumination microscopy (SIM) images revealed colocalization of Rab7-CLCA1 with lysosomes (Fig 7I). Further, Rab7 was knocked down in HT29-MTX-E12 cells and the process of CLCA1 fusion with lysosomes was imaged using confocal microscopy. The images revealed less colocalization of CLCA1 with lysosomes upon Rab7 knock-down as compared to vehicle control (Fig 7J and Fig 7K). Together, these data suggested modulation of CLCA1 protein degradation by Rab7 in goblet cells. Overall, our data highlights a crucial role of Rab7 in maintaining gut homeostasis. Rab7 downregulation, as observed during colitis, results in increased CLCA1 in goblet cell and thereby a higher secretion. These changes adversely affect the mucus layer, the composition of microbiota and epithelial barrier function altogether leading to inflammation (Fig 8).

**Fig 8.**
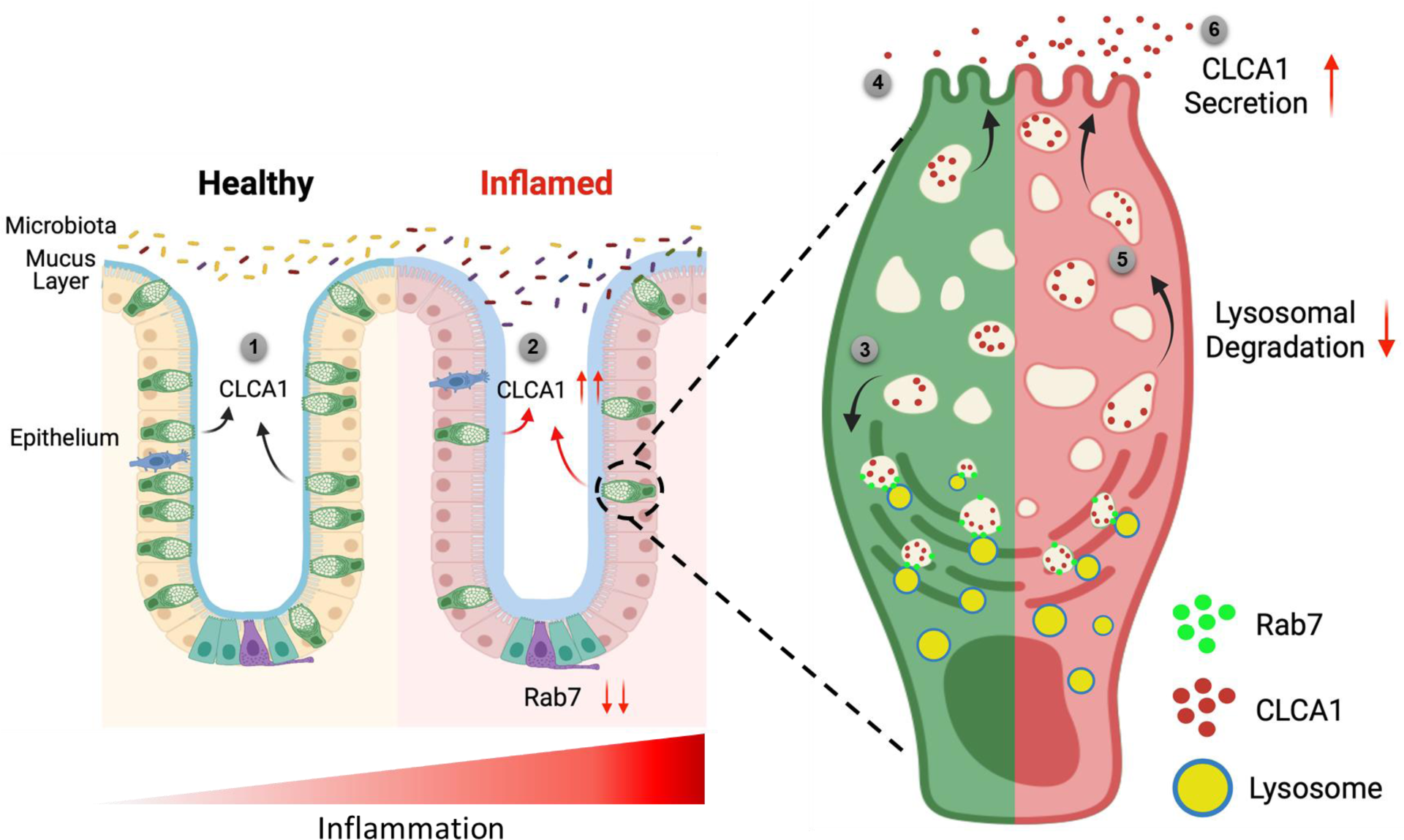
Rab7 maintains mucus layer dynamics in intestine by regulating degradation of CLCA1 protein via lysosomal fusion. Healthy intestine inhabits lumen microbes well separated by mucus layer secreted by goblet cells along with CLCA1 protein in balanced levels (1). During colitis, Rab7 downregulates along with increased expression of CLCA1 resulting in diffused mucus layer penetrable to microbes (2). In a goblet cell, CLCA1 filled vacuoles destined for secretion are rerouted for degradation pathway by Rab7 and fuse with lysosomes (3) leading to a balanced release outside the cell (4). However, during inflammation the loss of Rab7 consequently impedes CLCA1 degradation (5) fostering its increased secretion from the cell (6) (Created with BioRender.com).

## Discussion

Here we report significant contribution of one of the essential cellular protein i.e. Rab7 GTPase in colitis pathogenesis. Rab GTPases are amongst the fundamental proteins of a cell regulating important cellular mechanisms like membrane trafficking (Hutagalung & Novick, 2011). A number of comprehensive studies summarizing the molecular, physiological and pathological aspects of vesicle transport system link trafficking proteins dysfunction with human diseases such as carcinoma, neurodegenerative disorders, charcot-marie-tooth type B and diabetes (BasuRay et al., 2013; Mafakheri et al., 2018; Millecamps & Julien, 2013; Tzeng & Wang, 2016). Interestingly, the involvement of Rab GTPases in underlying mechanisms implicating IBD is still not well understood. Rab5, Rab34 and Rab11 have earlier been reported to contribute to correct assembly and maintenance of junctional complexes in intestinal epithelial cells (Citalán-Madrid et al., 2013), (Talmon et al., 2012). Dislocation of Rab13 has been shown in patients with Crohn’s disease (Oshitani, 2009).

Rab7 protein was strikingly observed to be down expressed during human and murine colitis. To explain the discordance about Rab7 getting upregulated in crypts of IBD patients as observed by Gang Du et al., 2020 (Du et al., 2020), we systematically analyzed different phases of inflammation development at diverse regions of intestinal mucosa. It seems Rab7 downregulation is dependent on stages of inflammation progression and the pattern varies between regions of the intestine viz. epithelium and crypts.

Using an inhibitor (CID1067700) of Rab7, its role in impairing B-cell class switching during murine lupus has been reported (Lam et al., 2016). Though specific to inhibit Rab7 GTPase activity, using this inhibitor limits the user to distinctively target any specific organ. In the current study we have exploited a recently created technique to deliver nucleic acids to intestine. Although being transient, this mouse model appears promising to briefly study and understand the outcomes of gut-specific knock-down of Rab7. Successful knock-down of Rab7 in mice intestine revealed several cues regarding its role in intestinal homeostasis. DSS treatment to Rab7^KD^ mice displayed worsening of inflammation in the intestine. This explains that down expression of Rab7 during colitis can be the cause of disease.

Ulcerative colitis pathogenesis is majorly associated with decreased number of goblet cells which in turn attributes to compromised mucus layer. A recent report describes that the reduction in mucus layer is related to reduced secretory function of remaining goblet cells (Birchenough et al., 2015). Though ubiquitously expressed in all cell types, dominant expression of Rab7 in goblet cells of intestine was unpredicted. The fact that Rab7 was specifically getting altered in goblet cells during colitis was rather intriguing and necessitated us to investigate in this aspect. Our findings revealed alterations in thickness and permeability leading to disrupted sterility in the inner mucus layer of Rab7^KD^ mice. Further, dysbiosis in luminal microbiota was also evident through 16S metagenomics analysis. Reduction in Firmicutes is the most consistent observation in IBD patients. Our data shows significantly decreased abundance of Clostridia, a class of Firmicutes in Rab7^KD^ mice.

Alterations in the mucus layer characteristics could be attributed to changes in its composition. A report on alterations of colonic mucus composition in UC shows that the structural weakening of mucus layer is due to reduction in 9 proteins among the 29 mucus core proteins such as Muc2, FCGBP, ZG16 and CLCA1. Mass spectrometry analysis revealed changes in the mucus composition upon Rab7 perturbation in mice colon. It was seen that CLCA1, among the few altered proteins, was surprisingly upregulated in Rab7^KD^ mice. Several omics studies in human UC patients have suggested variations in the CLCA1 protein expression during inflammation (Massimino et al., 2021). In an attempt to investigate this protein in colitis, a series of studies negated the involvement of CLCA1 in gut inflammation until recently by using *ex vivo* assays it was revealed that CLCA1 is important in maintaining the intestinal homeostasis and possess Muc2 cleaving properties thus plays role in regulating the mucus dynamics. CLCA1 poses to be an apt candidate to justify modifications in mucus layer upon Rab7 knock-down in mice.

Lack of knowledge regarding the secretion, regulation and stability of CLCA1 in a cell led us to investigate it at a deeper level. Our data reveals that Rab7 is responsible to direct the CLCA1 containing vesicles towards lysosomes for further degradation. No report has shown the presence of its free molecules in cytoplasm. However, we are not certain whether CLCA1 exclusively localizes in vesicles. It is evident that goblet cells maintain a basal level of CLCA1 to be secreted out in order to maintain the mucus dynamics and homeostasis. Rab7 perturbation disturbs this balance and thus leads to disrupted mucus layer dynamics and sterility which may initiate inflammation. Together, in the present study we establish the importance of Rab7 in maintaining the intestinal homeostasis by regulating the secretory functioning in the gut. Specifically, the involvement of Rab7 in managing core mucus component CLCA1 opens up avenues for therapeutic interventions. Our findings demonstrate the importance of a fundamental protein like Rab7 which can pose to be the root causative for initiating incurable diseases like IBD.

## Materials and Methods

### Human

Human samples were collected from the All India Institute of Medical Sciences (AIIMS), New Delhi for Non-IBD (IBD suspects), UC and CD groups with age over 18 and below 60 years with inclusion and exclusion criteria specified in Salman et. al., 2017. Two to four biopsies from each patient were collected to be used for qRT-PCR, western blotting and sectioning. Informed consent forms were acquired from all the patients. Influence of gender was not considered in the present study. UC severity was determined on the basis of ulcerative colitis endoscopic index of severity (UCEIS) score. Score 0-1 were considered under remission, 2-4 as mild and 5-6 as moderate. Ethics approval for the use of human samples were obtained from both the institutes Regional Centre for Biotechnology (RCB) and AIIMS.

### Mice

C57BL/6 mice were bred and housed in pathogen-free conditions, provided sterilized food and water at 25℃ with 12-h light/dark cycle in Small Animal Facility of RCB, Faridabad. For DSS colitis experiments, female C57BL/6 mice 6-8 week old, weighing 18-20gms were used. Mice were fed with 2.5% DSS (w/v) (36–50 kDa molecular weight; MP Biomedical) for 3, 5 and 7 days in autoclaved water. Autoclaved water was given to control mice. Body weights were monitored daily along with colitis symptoms like rectal bleeding and diarrhea. Colon, spleen, liver and MLN were harvested. Colon and spleen were examined for changes in length and size. Organs were further used for western blots. Distal colons were fixed with 4% paraformaldehyde and processed for sectioning.

### Cell culture and transfection

HT29 (Lot. 09K003) and HT29-MTX-E12 (Lot. 18K206) cell lines were obtained from ECACC. HT29 cells were cultured in RPMI (Merck) containing 10% fetal bovine serum (v/v) (Gibco), 1% sodium pyruvate (Gibco), 2gm/litre sodium carbonate (Sigma) and 1% penicillin/streptomycin (Gibco). For differentiating into mucin secreting goblet-like cells, HT29 cells were grown in glucose free DMEM (Gibco), supplemented with galactose (250mM) for 3 days. HT29-MTX-E12 cells were grown in DMEM glutamax medium (Thermo Fisher Scientific) supplemented with 10% fetal bovine serum (v/v), 1% sodium pyruvate and 1% penicillin/streptomycin. Plasmid transfections and siRNA mediated knock downs were done along with cell seeding using Transfectin reagent (BioRad) and Dharmafect reagent (Horizon discovery) respectively. To inhibit lysosomal and proteasomal degradation, cells were treated with 100nm Bafilomycin (Sigma) and 10µM MG132 (Sigma) for 6 hours and 8 hours respectively.

### Intestinal Epithelial Cells and crypt isolation

Mice colons were flushed with ice-cold PBS to remove luminal content and were opened longitudinally using sterile scissors. Mucus was removed gently using a soft rubber scraper. Colons were treated with 30mM EDTA for 30 minutes at 4℃. Cells were scrapped from the colon surface in ice-cold PBS and centrifuged at 2000 rpm for 5 minutes to collect intestinal epithelial cells for lysate preparation (Mustfa et al., 2017). For colonic crypt isolation, intestinal epithelial organoid culture kit protocol (Stem cell technologies) was followed as per the manufacturer’s instructions. Briefly, colons were washed with DPBS (without Ca^2+^ and Mg^2+^) and minced into small pieces. After thorough washing, colon pieces were treated with gentle cell dissociation reagent. Fractions of crypts from solution were collected in 0.1% BSA solution after passing through 70µm strainer. Crypts were further lysed in RIPA buffer for western blotting.

### CRISPR-Cas9 mediated Rab7 knockout in HEK293T cells

Rab7 knockouts were generated in HEK293T cells using CRISPR-Cas9 technique. sgRNA sequence for human Rab7a was selected (AGGCGTTCCAGACGATTGCA) with AGG as protospacer region (NGG). The gRNA sequence was further cloned into cas9 vector pSpCas9(BB)-2A-Puro (PX459) from addgene. The clone was confirmed by sequencing. 1.3×10^5^ HEK293T cells were seeded and cultured for 24 hours before transfection with Rab7-knockout plasmid using Lipofectamine 2000 (Thermo). Clonal selection of cells was done further by serial dilution as 0.5 cells/well of 96-well plate. Each clone was then tested for Rab7 deletion by western blotting with anti-Rab7 antibody (Sigma).

### Quantitative RT-PCR

Total RNA from the human biopsy samples and mice colon tissue was isolated using RNeasy mini kit (Qiagen) according to the manufacturer’s protocol. cDNA was prepared using 1µg of total RNA for each sample using cDNA synthesis kit (Bio-Rad). Real time PCR was performed utilizing SYBR green master mix (BioRad) for 20µl reaction volume in 96-well plates in CFX96 Real-Time system (BioRad). 18S and HPRT genes were used as housekeeping to normalize the reactions according to the sample origin. The list of primers is included in Table S2.

### Western blotting

Human biopsies and mice organ tissues were homogenized using tissue homogenizer (Precellys) in RIPA buffer with added protease arrest (G Biosciences) to prepare protein lysates. Cells were lysed in RIPA buffer (with protease arrest) after PBS wash. Protein amount was measured by BCA solution (Sigma). Equal amount of protein samples was separated on SDS-PAGE gel (12%) electrophoresis and transferred to nitrocellulose membrane (BioRad). Blots were blocked with 5% skim milk for 1 hour at room temperature and probed with antibodies at 4℃ overnight against the desired proteins. The primary antibodies used were anti-Rab7 (Sigma, R4779, 1:5000), anti-GAPDH (Invitrogen, 39-8600, 1:2000), anti-β-Actin (Cell Signaling Technology, 4970S, 1:20,000), anti-CLCA1 (Abcam, ab180851, 1:20,000). Specific secondary antibodies conjugated with HRP were probed for 1 hour at room temperature. Blots were detected and visualized for protein bands with Immobilon Forte western HRP substrate (Millipore) and imaged in Image Quant LAS4000. Band intensities were measured by ImageJ software.

### Histology and immunostaining

Mice distal colons were sectioned (5µm) after fixing in 4% paraformaldehyde. For histopathology analysis, sections were stained with hematoxylin and counter stained with eosin. Histological scoring of colon sections was evaluated by a blinded pathologist following these parameters: loss of epithelium (0-3), crypt damage (0-3), depletion of goblet cells (0-3) and inflammatory cell infiltrate (0-3). For immunostaining, human biopsy and mice colon samples were sectioned and after fixing with 4% paraformaldehyde antigen retrieval was performed. The sections were probed with anti-Rab7 (Sigma, R4779, 1:400), UEA1-FITC (Vector Laboratories, FL-1061, 1:400), anti-Muc2 (Santa Cruz, sc-515032, 1:200), anti-CLCA1 (Abcam, ab180851, 1:100) in blocking (5% goat serum) overnight. Sections were incubated with HRP or fluorophore tagged secondary antibodies for 2 hours. Nucleic acid was stained with DAPI (1µg/ml). Sections were cured and mounted with Prolong Gold Antifade Reagent (Thermo Fisher). Sections were imaged in confocal microscope (Leica SP8).

### Transmission and Immuno Electron Microscopy

Ultrastructure of HT29-MTX-E12 cells was visualized by Transmission Electron Microscopy (TEM). Cells grown in complete medium were prefixed with Karnovsky fixative (2.5% Glutaraldehyde + 2% paraformaldehyde in 0.1M phosphate buffer, pH 7.4). Post fixation with 1% osmium tetroxide in 0.1M phosphate buffer for 1 hour, cells were dehydrated in a graded series of ethanol solutions: 30%, 50%, 70%, 80%, 90% and 100% and with acetone and toluene. Samples were further embedded in Epon 812 resin and polymerized at 65℃ for 48 hours. Ultrathin sections of 70nm were cut, followed by staining with uranyl acetate and lead acetate and mounted on a grid. Imaging was done with TALOS 200S Transmission electron microscope (Thermo Fisher Scientific) operated at 200 kV. For Immuno Electron Microscopy (IEM), cells were fixed in 0.5% glutaraldehyde in 0.1M phosphate buffer and were embedded in LR white resin. Thin sections (70nm) were prepared and incubated for 30 minutes at room temperature in 1% BSA, followed by overnight incubation with rabbit anti-Rab7 antibody (1:1000 in PBS with 1% BSA) in a humid chamber at 4℃. After washing with 1% BSA, sections were incubated with 18nm gold affinity pure goat anti-rabbit IgG antibody (Jackson ImmunoResearch laboratories, 1:50 in 1% BSA) for 2 hours at room temperature. Sections were then contrasted and examined as described above.

### Alcian blue and Fluorescence In-situ Hybridization staining

Mice distal colon pieces containing fecal pellet were fixed in Carnoy’s fixative (60% methanol+30% chloroform+10% glacial acetic acid), embedded in paraffin and sectioned (5µm). Sections were dewaxed using Histoclear (Sigma), stained with alcian blue solution and counter stained with nuclear fast red (Musch et al., 2013). Sections were visualized in Nikon bright field microscope and mucus thickness was measured using ImageJ software. For Fluorescence in-situ hybridization (FISH), dewaxed sections were incubated with 1.3µg of general bacterial probe EUB338-Alexa Fluor 647 overnight at 50℃ in a humid chamber (Swidsinski, 2005). Nucleus was stained with DAPI (1µg/ml). Sections were imaged in confocal microscope (Leica SP8). Bacteria in mucus layer per section was calculated using particle counter in ImageJ software.

### Immunocytochemistry and Structured Illumination Microscopy

Cells were seeded on 18mm glass coverslips were fixed with 4% methanol free paraformaldehyde and blocked in 0.1% BSA+0.01% TritonX-100 for 1 hour. Cells were probed with anti-CLCA1 (Abcam, ab180851, 1:100), anti-Muc2 (Santa Cruz, sc-515032, 1:200), anti-Rab7 (Sigma, R4779, 1:400) and Phalloidin-594 () at 4℃ overnight. Cells were further incubated with fluorophore tagged secondary antibodies for 2 hours. Nucleic acid was stained with DAPI (1µg/ml). Coverslips were mounted with Prolong Gold Antifade Reagent (Thermo Fisher) and visualized in confocal microscope (Leica SP8) for immunocytochemistry and in Elyra PS1 (Carl Zeiss) for structured illumination microscopy.

### Polymer based knock-down in mice

Female C57BL/6 mice (aged 6-8 weeks, weight 18-23gms) were divided into six groups. Group 1 was untreated control group. Group 2 mice were treated with control nanogels only. Group 3 mice were treated with Rab7 siRNA nanogels. Group 4 mice were fed with only ∼2.5% DSS. Group 5 mice were fed with ∼2.5% DSS and treated with control nanogels. Group 6 mice were fed with ∼2.5% DSS and treated with Rab7 siRNA-nanogels. For one mouse, Rab7 siRNA nanogels were prepared by incubating TAC6 (100μL of stock 2 mg/mL) with siRNA (0.6 μg/per dose) for 30 min., followed by incubation with 10μL of SPA (2 mg/mL) for 10 min. For control nanogels, scrambled siRNA was fed.

### Mucus Isolation and separation

Mice colons were harvested from each mouse and after washing with ice-cold PBS were cut open longitudinally. Mucus was scraped using rubber scraper, resuspended in PBS+Protease inhibitor cocktail (2X) and snap frozen. Further, samples were denatured in reducing buffer at 95℃ for 20 minutes followed by additional reduction at 37℃ for 1 hour. Composite gradient Agarose-PAGE gel was prepared with 0.5-1% Agarose, 0-6% Polyacrylamide and 0-10% Glycerol and solidified at 37℃ for 1 hour followed by overnight incubation in a humid chamber. Mucus samples were separated on mucin gel electrophoresis and stained with Alcian blue to visualize highly glycosylated mucins (Johansson et al., 2009).

For isolation of mucus from human samples, the biopsy tissue was treated with N-acetyl cysteine (2% PBS solution) for 5 minutes at 4℃. The supernatant was collected and centrifuged at 2000 rpm for 5 minutes to remove cell debris. Protease inhibitor (2X) was added to the supernatant. The mucus samples were further denatured in reducing buffer at 95℃ for 15 minutes.

### Mucus proteomics by LC MS/MS

LC-MS/MS analysis was performed using mucus sample lysates. In-Gel digestion was carried out using Trypsin gold (Promega). The samples were purified using C18 SepPak columns (Thermo, USA). The peptide samples were dissolved in 98% milliQ-H_2_O, 2% acetonitrile, 0.1% formic acid. Tandem MS analysis was performed using a 5600 TripleTOF analyser (ABSCIEX) in Information Dependent Mode. Protein identification was performed with MaxQuant software (v1.6.0.16) under default settings, except for the peptide length that was set from 6 to 40. Masses were searched against the Mouse Uniprot Proteome (downloaded 27/4/20) with additional mice mucin database, VerSeDa, antimicrobial peptides (from uniport) and cytokines (from uniprot). The peptides resulting from MaxQuant were then processed as follows: Known MS contaminants and reverse sequences were filtered out; we allowed peptides that had at least two valid LFQ intensities out of sample replicates, and we included razor peptides, which belong to a unique MaxQuant ‘protein group’; finally, missing values were imputed to a random value selected from a normal distribution of 0.3 s.d. and downshifted 1.8 s.d. The data processing was performed in python v3.8. Statistical analyses were performed in R v4.0.2. In R data was visualized using the complexheatmap and ggplot packages (Gu et al., 2016; Wickham, 2009). Protein annotation and gene ontology analysis were performed with the gene-set enrichment analysis using the enrichR webtool (Chen et al., 2013; Kuleshov et al., 2016; Xie et al., 2021).

### 16S Metagenomic profiling

Fecal pellets were collected from each mouse and submitted to miBiome Therapeutics for processing and analysis. Bacterial genomic DNA was isolated by QIAamp PowerFecal Pro DNA Kit (Qiagen, 51804). DNA samples were quality checked and subjected to library preparation in alignment with the 16S metagenomic library preparation protocol from Illumina Inc. to target the 16S V3 and V4 region. The full-length primer sequences targeting the V3-V4 region are: 16S Amplicon PCR Forward Primer 5’TCGTCGGCAGCGTCAGATGTGTATAAGAGACAGCCTACGGGNGGCWGCAG and 16S Amplicon PCR Reverse Primer 5’GTCTCGTGGGCTCGGAGATGTGTATAAGAGACAGGACTACHVGGGTATCTAAC. Libraries were diluted to 4 nM, pooled, spiked with 20% PhiX pre-made library from Illumina and loaded on a MiSeq v3 kit. Sequencing was performed for 2×300 cycles. The original raw data obtained from Illumina platform are recorded in FASTQ files, which contains sequence information (paired end reads) and corresponding sequencing quality information (Q score). The reads obtained from the instrument were made adapter free, using the adapter trimming of local run manager on Illumina. The paired-end reads were assembled, filtered, trimmed and aligned to SILVA 16S database using mother software v.1.44.1. The reads were clustered based on similarity, and further clustered into OTUs (Operational Taxonomic Unit) using mothur software v.1.44.1 at 97% identity against database greengenes_13_8_99.

### Enzyme Linked Immunosorbent Assay

Colonic mucus was scrapped and diluted in PBS. The homogenate was centrifuged to remove debris and the supernatant was transferred to a new tube. The samples were quantified by BCA assay and further equal protein was assayed by enzyme-linked immunosorbent assay (ELISA) for TNF-a using the manufacturer’s protocol (R&D systems, USA).

### Statistical Analysis

Results were analyzed and plotted using GraphPad prism 8.0.1 software. All results were expressed as mean standard error from individual experiment done in triplicates. Data were analyzed with standard unpaired two-tailed Student’s t test and the Welch’s t test where applicable. p values less than 0.05 were considered as statistically significant.

## Supplementary Materials

Fig S1. Rab7 expression in various states of murine and human colitis (related to Fig 1)

Fig S2. Expression of Rab7 in goblet cells of different types and regions of gastrointestinal tract (related to Fig 2)

Fig S3. Rab7 perturbation changes goblet cell number and Muc2 distribution (related to Fig 3 and Fig 4)

Fig S4. Gut microbiota composition analysis of Rab7^KD^ mice using 16 S rRNA sequencing (related to Fig 5)

Fig S5. Mucus proteome analysis of Rab7^KD^ mice (related to Fig 6)

Fig S6. CLCA1 protein degrades via lysosomal degradation pathway (related to Fig 7)

Table S1. List of patient clinical parameters

Table S2. List of primers used in qRT-PCR

## Acknowledgements

We thank the Mass spectrometry facility, the Central Instrumentation Facility (CIF) of Regional Centre for Biotechnology (RCB) and Small Animal facility at NCR, Biotech Science Cluster. We also thank Sophisticated Analytical Instrumentation Facility, AIIMS (SAIF-AIIMS) and Advanced Technology Platform Centre (ATPC) for imaging facilities. We are thankful to Professor Gunnar C Hansson for Muc2 plasmids and Dr. Amit Tuli for Rab7 plasmids. We acknowledge Addgene for the plasmid. We thank Dr. Bobby Cherayil for all the discussions and feedback during the manuscript preparation.

## Funding

This work was supported by DBT grant (BT/PR45284/CMD/150/9/2022), RCB Core Grant, ICMR-SRF award of PG, CSIR-JRF programme (Council of Scientific and Industrial Research, Govt. of India) of PG.

## Author contributions

Conceptualization: C.V.S.

Methodology: P.G., C.V.S.

Investigation: P.G., Y.R., B.S.

Formal Analysis: V.F.Y

Resources: M.M., S.C., S.T.

Supervision: G.M., A.S, S.C.Y., A.K.P., Y.M, A.B, V.A.

Writing—original draft: P.G., C.V.S.

Writing—review & editing: P.G., C.V.S., G.M., Y.M.

### Competing Interests

The authors declare that they have no competing interests.

### Ethics

Animals ethics proposal was approved by the RCB Institutional Animal Ethics Committee (approval no. RCB/IAEC/2020/083). Informed consent form was obtained from all the patients and the study protocol was submitted to the ethics committee of the institutes: RCB (IEC-141/05.03.2021, RP-34/2021) and AIIMS (IEC/NP-189/2013&RP-12/17.06.201307.06.2013).

## Notes

### Competing Interest Statement

The authors have declared no competing interest.

### Summary of Updates

Edits in main text. Figure S2C and Figure S2D updated. Author affiliation updated.

