## Supplementary Information for "Rab7 dependent regulation of goblet cell protein CLCA1 modulates gastrointestinal homeostasis"

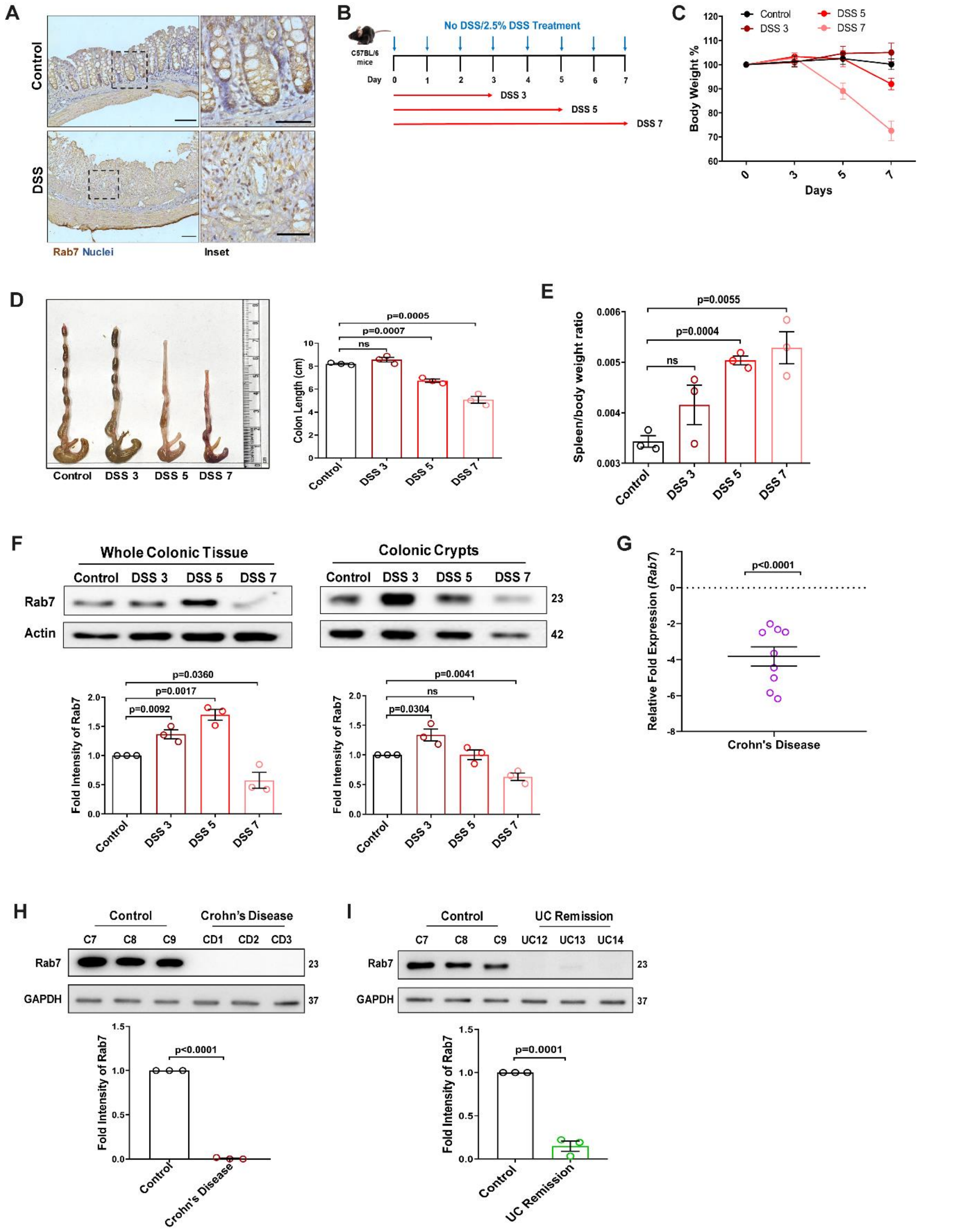


**Fig S1. Rab7 expression in various states of murine and human colitis (related to Fig 1)**

(A) Immunohistochemistry images of Rab7 staining (brown color) in distal colon sections of healthy and DSS-treated mice (Scale bar=100µm). Inset shows zoomed areas of the image (Scale bar=50µm).

(B) Schematic representation of DSS treatment to mice for different time durations.

(C) Graph represents body weight percent of mice in different groups.
(D) Gross morphology of colon and caeca. Graph shows colon length quantification.
(E) Spleen to body weight ratio indicating increased splenomegaly and inflammation.
(F) Dynamics of Rab7 expression in whole tissue and crypts isolated from the intestines of healthy and DSS-treated mice for different time durations. Corresponding graph shows densitometry analysis of Rab7 expression normalized to loading control (β actin).
(G) A pilot study of Rab7 expression in CD patients. RT-PCR analysis of relative fold expression of *Rab7* gene in human CD patient colonic biopsies (n=9) relative to average control values (n=10). *HPRT* was used for normalization.
(H) Immunoblotting of Rab7 protein in CD (n=3) and control (n=3) biopsy samples. GAPDH was used as loading control.
(I) Immunoblotting of Rab7 protein in human UC remission (n=3) and control (n=3) biopsy samples. GAPDH was used as loading control.
Each dot represents (D, E) one mouse or (F, G and H) one human. Error bars represent mean+SEM. Statistical analysis by Student's t test. ns=non-significant.

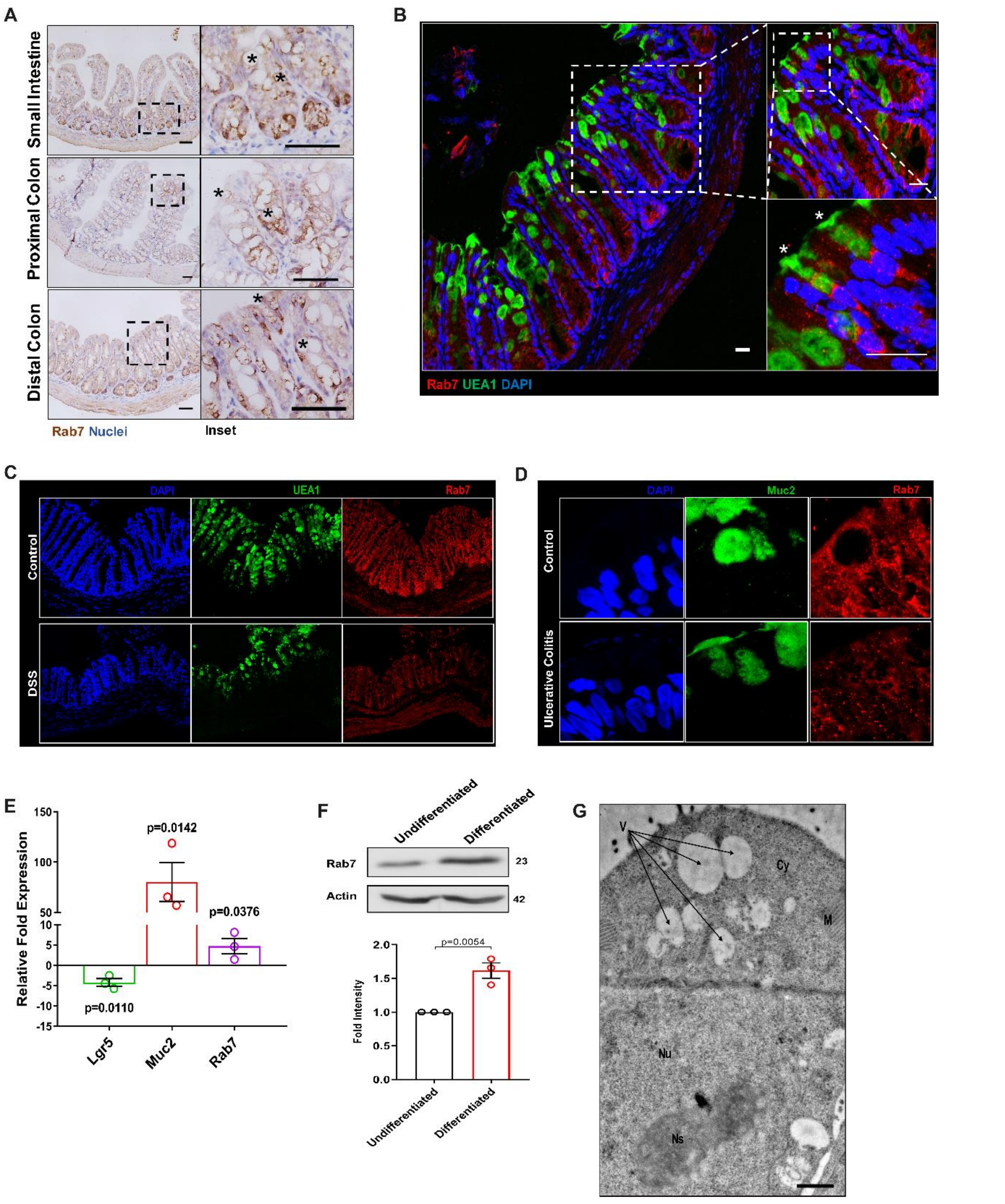


**Fig S2. Expression of Rab7 in goblet cells of different types and regions of gastrointestinal tract (related to Fig 2)**

(A) Immunohistochemistry images of staining of Rab7 in different regions of intestine: small intestine, proximal colon and distal colon; showing Rab7 expression in vacuolated cells (marked with asterisk) (n=3). Scale bar=50µm.

(B) Confocal images of mice distal colon stained with Rab7 (red) and goblet cell specific marker UEA1 (green) showing presence of Rab7 in UEA1 positive cells (marked with asterisk) (n=3). Scale bar=100µm.

(C-D) Single-channel images of DAPI, UEA1/Muc2 and Rab7 staining shown in Fig. 2A and Fig. 2C.
(E) RT-PCR analysis of relative fold expression of *Lgr5*, *Muc2* and *Rab7* genes in differentiated HT29 cells relative to undifferentiated cells. *HPRT* was used for normalization.
(F) Immunoblot showing Rab7 protein expression in differentiated HT29 cells relative to undifferentiated cells. Graph represents densitometric analysis of Rab7 expression calculated by normalizing to loading control (β actin).
(G) TEM analysis of HT29-MTX-E12 cells showing presence of vacuoles (V) in the cytoplasm (Cy) along with nucleus (Nu) nucleolus (Ns) and mitochondrion (M). Each dot represents one individual experiment. Error bars represent mean+SEM. Statistical analysis by Student's t test. ns=non-significant.

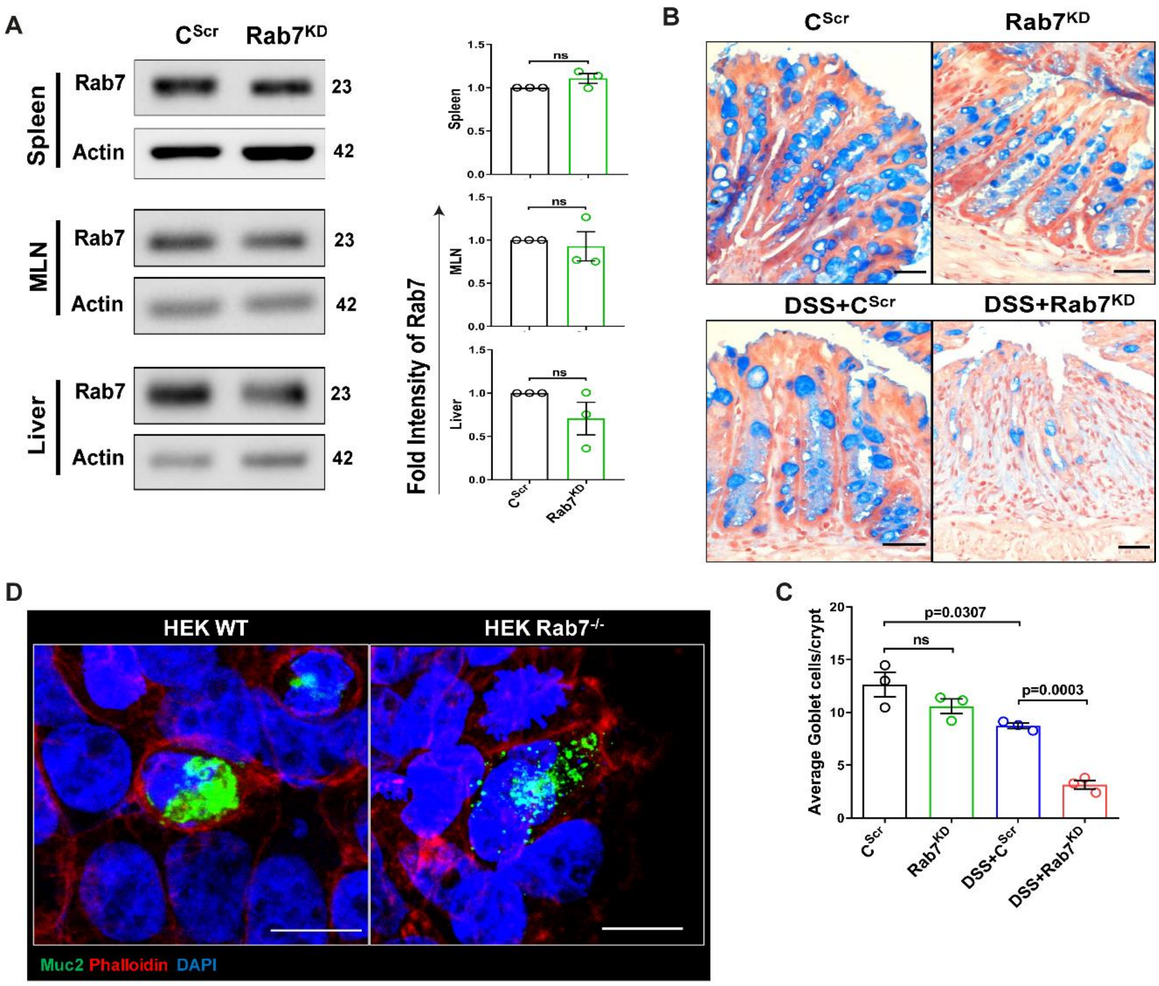


**Fig S3. Rab7 perturbation changes goblet cell number and Muc2 distribution (related to Fig 3 and Fig 4)**

(A) Rab7 protein expression in lysates of spleen, MLN and liver collected from $C^{Scr}$ and $Rab7^{KD}$ mice. Graph represents densitometric analysis showing fold intensity of Rab7 expression calculated by normalizing to loading control (β actin).
(B-C) Alcian blue staining in sections of distal colon show mucus filled goblet cells (blue) and nuclei (red). Number of goblet cells per crypt were counted manually (15-20 crypts per mouse).

(D) Representative immunofluorescence images of HEK WT and Rab7$^{-/-}$ cells stained for Muc2 (green), Phalloidin (red) and cell nuclei (blue) (Scale bar=10μm).
Each dot represents one mouse. Error bars represent mean+SEM. Statistical analysis by Student's t test. ns=non-significant.

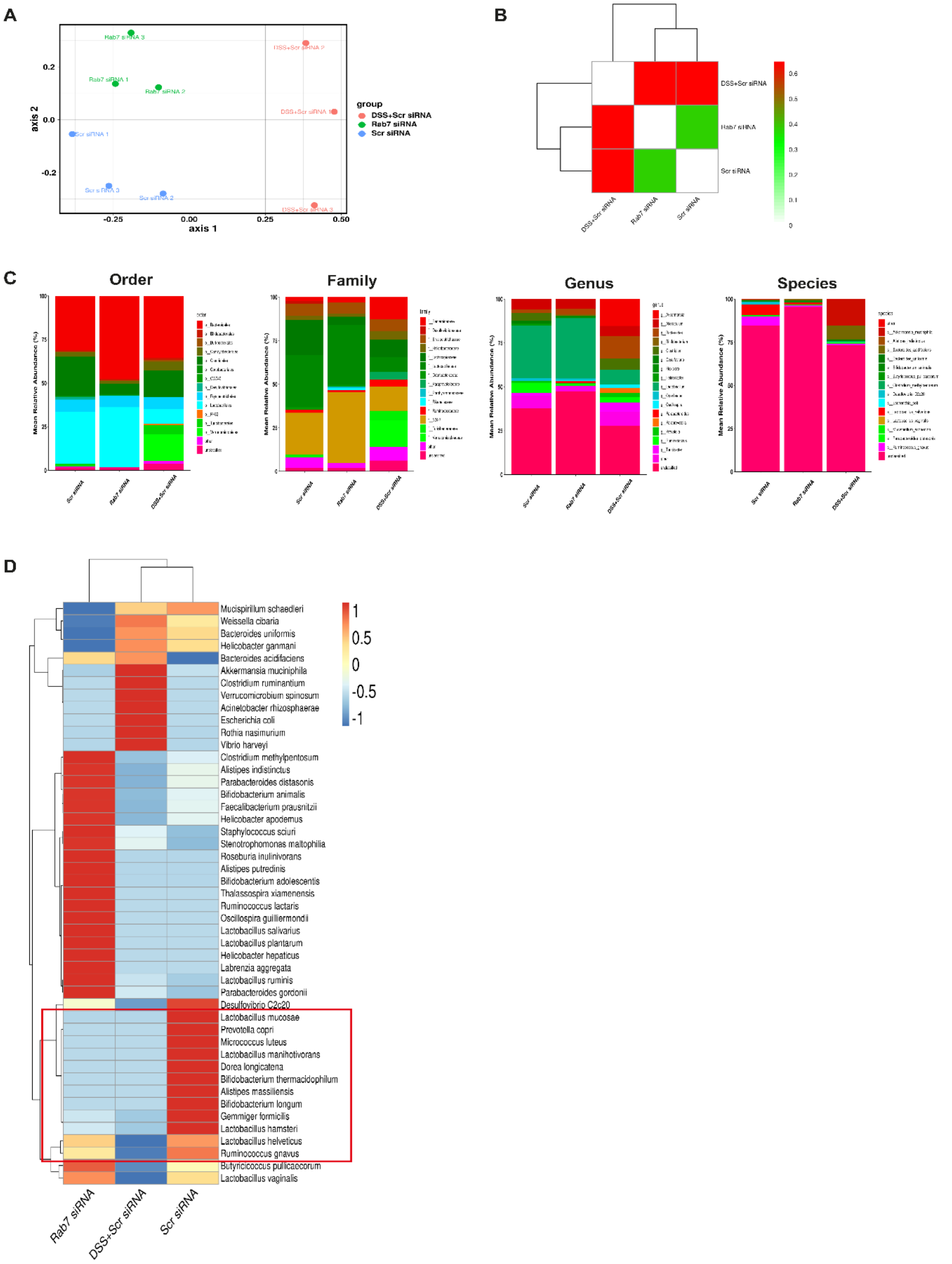


**Fig S4. Gut microbiota composition analysis of Rab7$^{KD}$ mice using 16 S rRNA sequencing (related to Fig 5)**

(A) Non-metric dimensional scaling (NMDS) calculated from distance matrices obtained from Bray-Curtis.
(B) Heat map showing beta diversity index obtained from Bray-Curtis.
(C) Mean relative abundance of top 10 Order, Family, Genus and species.
(D) Heat map representing mean relative abundance of all species across different groups (blue represents low abundance, red represents more abundance).

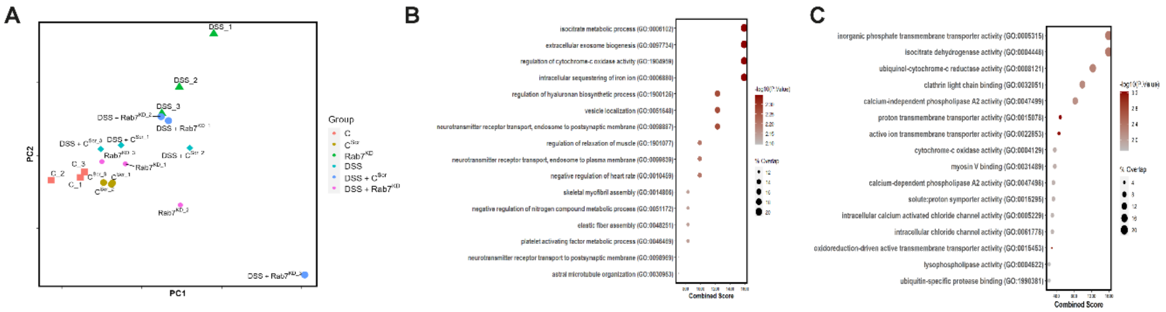


D

**Rab7$^{KD}$ vs C$^{Scr}$**

| Gene Name | Cellular Location | p value | Log2 Fold Change |
|---|---|---|---|
| Hist1h1d | Nucleus | 0.033024 | 3.159073 |
| Hint1 | Cytoplasm, Nucleus | 0.024104 | 2.04931 |
| Ndufa4 | Mitochondria | 0.042742 | 1.954867 |
| Pla2g4c | ER, Mitochondria | 0.017834 | 2.307447 |
| Sri | Cytoplasm | 0.047669 | 1.54052 |
| Nmes1 | Nucleus | 0.001065 | 1.831148 |
| Slc25a3 | Mitochondria | 0.003168 | 1.745184 |
| Clca1 | Secreted | 0.003566 | 2.334244 |
| Pcbp1 | Cytoplasm, Nucleus | 0.001547 | -2.44554 |
| Idh3a | Mitochondria | 0.028413 | -2.15885 |
| Ppa1 | Cytoplasm | 0.000799 | -1.70347 |

E

**DSS+Rab7$^{KD}$ vs DSS+C$^{Scr}$**

| Gene Name | Cellular Location | p value | Log2 Fold Change |
|---|---|---|---|
| Eef1b | Cytoplasm | 0.036837 | 1.997831 |
| Alb | Secreted | 0.035046 | 1.734723 |
| Apoa1 | Secreted | 0.01182 | 2.105002 |
| Gdi2 | Cytoplasm | 0.010669 | 2.118751 |
| Atp1a1 | Cell Membrane | 0.011803 | 1.593363 |
| Tf | Secreted | 0.010292 | 1.584418 |
| Clca1 | Secreted | 0.004133 | -2.56531 |
| Prdx5 | Mitochondria, Peroxisomes | 0.034956 | -2.25185 |

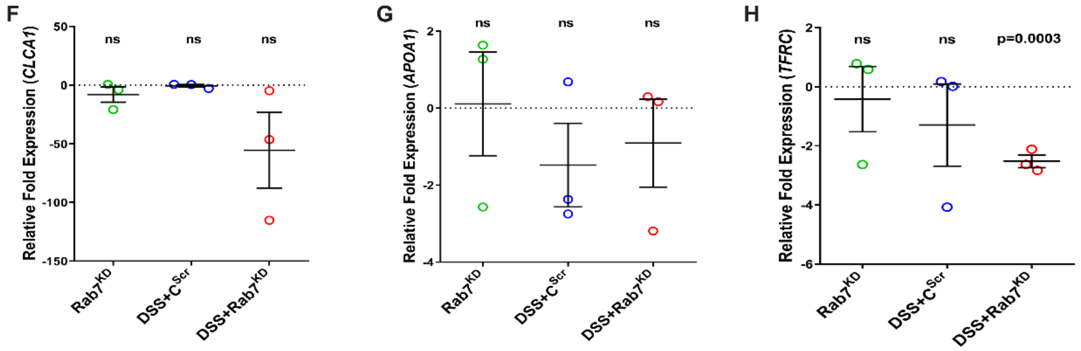


**Fig S5. Mucus proteome analysis of Rab7$^{KD}$ mice (related to Fig 6)**
(A) Principle component analysis (PCA) of different experimental groups.
(B-C) Fold change of proteins enriched in Rab7$^{KD}$ mice group compared to C$^{Scr}$ mice group based on Gene ontology analysis of biological process (B) and molecular function (C).
(D-E) Table summarizing fold change of differentially expressed proteins in Rab7$^{KD}$ versus C$^{Scr}$ mice group (D) and DSS+ Rab7$^{KD}$ versus DSS+C$^{Scr}$ treated mice group (E).

(F-H) Relative fold expression of *CLCA1* (F), *APOA1* (G) and *TFRC* (H) in Rab7$^{KD}$, DSS+C$^{Scr}$ and DSS+Rab7$^{KD}$ treated mice relative to C$^{Scr}$ mice (n=3). *HPRT* was used for normalization.
Each dot represents one mouse. Error bars represent mean+SEM. Statistical analysis by Student's t test. ns=non-significant.

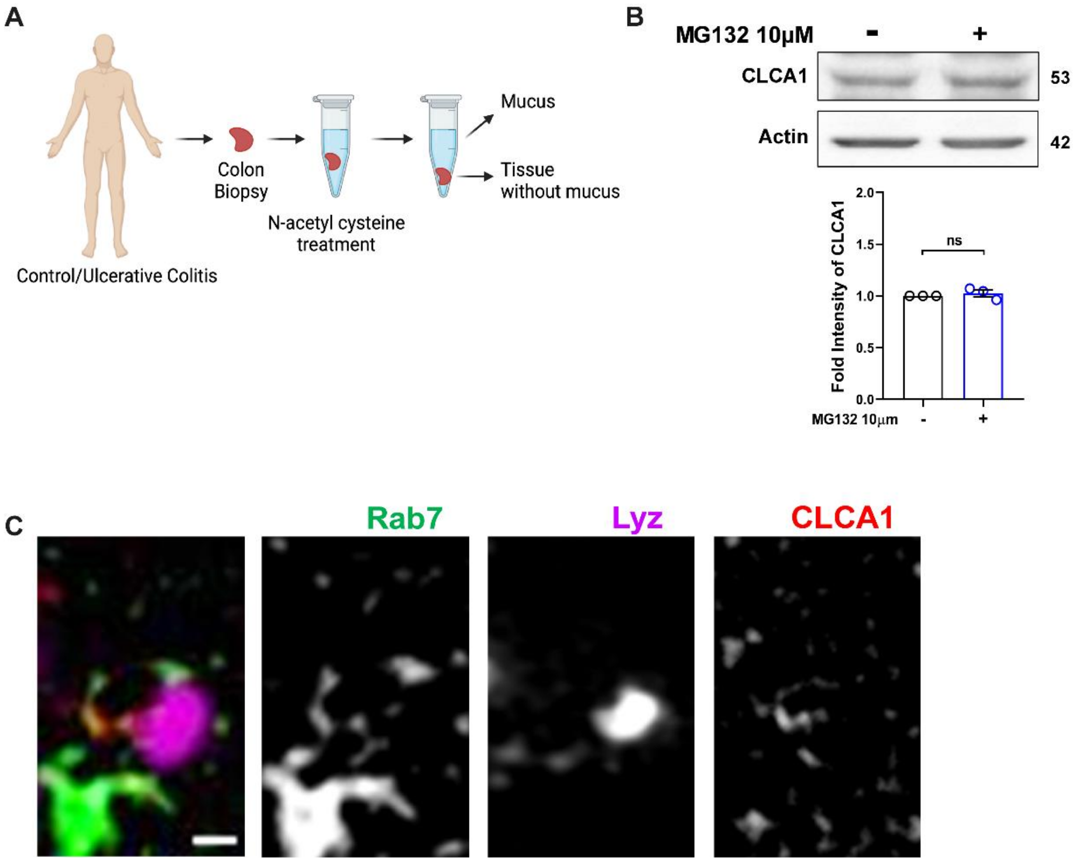


**Fig S6. CLCA1 protein degrades via lysosomal degradation pathway (related to Fig 7)**
(A) Schematic illustrating mucus isolation steps from human colon biopsy samples. (Created with BioRender.com)
(B) Immunoblot showing CLCA1 protein expression after MG132 treatment. Graph represents densitometric analysis showing fold intensity of CLCA1 expression calculated by normalizing to loading control (β actin).
(C) Representative zoomed in SIM images of HT29-MTX-E12 cells showing Rab7, CLCA1 and Lysosomes in grey scale. The left panel show the merged image.
Each dot represents one individual experiment. Error bars represent mean+SEM. Statistical analysis by Student's t test. ns=non-significant.

**Table S1.** List of patient clinical parameters

| Variables | Ulcerative Colitis | Crohn's Disease |
|---|---|---|
| Gender<br>Male<br>Female | <br>17<br>11 | <br>5<br>4 |
| Age in years (Mean±SD) | 35.28±10.54 | 34.11±14.85 |
| Duration of Disease in years (Mean±SD) | 5.13±5.15 | 5.77±2.9 |
| UCEIS (0-8)<br>Remission (0-1)<br>Mild (2-4)<br>Moderate (5-6) | <br>3<br>19<br>6 | NIL |
| Disease extent | Left-side colitis: 20<br>Pan colitis: 6 | Left-side colitis: 3<br>Colonic: 2<br>Ileocolonic: 3<br>Distal Ileal: 1 |
| Concomitant medications<br>5-aminosalicylic acid<br>Azathioprine<br>Steroids | <br>25<br>6<br>0 | <br>1<br>8<br>0 |

**Table S2.** List of primers used in qRT-PCR

| Gene Name | Forward Primer (Direction 5'-3') | Reverse Primer (Direction 5'-3') |
|---|---|---|
| Human Rab7 | CATCCTGGGAGATTCTGGAGTC | TGTGTCCCATATCTGCATTGTG |
| Human CLCA1 | TTTGTTCTCCAATCCCGCCA | CGGATCACTTCCCATGTGCT |
| Mouse CLCA1 | GAACAACAACGGCTATGAGGG | GCCTGAGTCACCATGTCCTT |
| Mouse APOA1 | ATTGACTCGGGACTTCTGGG | AATTCGTCCAGGTAGGGCTG |
| Mouse TFRC | TATCTTCTGGGGCTCTGGCT | CAGGGCCAACTGGTTTCTGA |
| Human Lgr5 | AAACCTCTCCAGCTGGGTAG | TTCAGCGATCGGAGGCTAAG |
| Human Muc2 | CGAAACCACGGCCACAACGT | GACCACGGCCCCGTTAAGCA |
| Human 18S | GAGGGACAAGTGGCGTTCA | CCGGACATCTAAGGGCATCA |
| Human HPRT | GCTATAAATTCTTTGCTGACCTGCTG | AATTAACTTTTATGTCCCCTGTTGACTGG |
| Mouse HPRT | CACAGGACTAGAACACCTGC | GCTGGTGAAAAGGACCTCT |
